# hibayes: An R Package to Fit Individual-Level, Summary-Level and Single-Step Bayesian Regression Models for Genomic Prediction and Genome-Wide Association Studies

**DOI:** 10.1101/2022.02.12.480230

**Authors:** Lilin Yin, Haohao Zhang, Xinyun Li, Shuhong Zhao, Xiaolei Liu

**Affiliations:** Key Laboratory of Agricultural Animal Genetics, Breeding and Reproduction, Ministry of Education, College of Animal Science and Technology, Huazhong Agricultural University, Wuhan, Hubei, 430070, PR China; School of Computer Science and Artificial Intelligence, Wuhan University of Technology, Wuhan, Hubei, 430070, PR China; Key Laboratory of Agricultural Animal Genetics, Breeding and Reproduction, Ministry of Education, Key Lab of Swine Genetics and Breeding of Ministry of Agriculture and Rural Affairs, The Cooperative Innovation Center for Sustainable Pig Production, College of Animal Science and Technology, Huazhong Agricultural University, Hubei Hongshan Laboratory Wuhan, Hubei, 430070, PR China; Key Laboratory of Agricultural Animal Genetics, Breeding and Reproduction, Ministry of Education, College of Animal Science and Technology, Shenzhen Institute of Nutrition and Health, Huazhong Agricultural University, Hubei Hongshan Laboratory, Wuhan, Hubei, 430070, PR China

**Keywords:** Genomic prediction, Genome-wide association studies, Bayesian regression, Single-step model, Summary statistics, hibayes, R

## Abstract

With the rapid development of sequencing technology, the costs of individual genotyping have been reduced dramatically, leading to genomic prediction and genome-wide association studies being widely promoted and used to predict the unknown phenotypes and to locate causal or candidate genes for animal and plant economic traits and, increasingly, for human diseases. Developing newly advanced statistical models to improve accuracy in predicting and locating for the traits with various genetic architectures has always been a hot topic in those two research domains. The Bayesian regression model (BRM) has played a crucial role in the past decade, and it has been used widely in relevant genetic analyses owing to its flexible model assumptions on the unknown genetic architecture of complex traits. To fully utilize the available data from either a self-designed experimental population or a public database, statistical geneticists have constantly extended the fitting capacity of BRM, and a series of new methodologies have been proposed for different application scenarios. Here we introduce **hibayes**, which is the only one tool that can implement three types of Bayesian regression models. With the richest methods achieved thus far, it covers almost all the functions involved in genetic analyses for genomic prediction and genome-wide association studies, potentially addressing a wide range of research problems, while retaining an easy-to-use and transparent interface. We believe that **hibayes** will facilitate the researches conducted by human geneticists, as well as plant and animal breeders. The **hibayes** package is freely available from CRAN at https://cran.r-project.org/package=hibayes.

## 1. Introduction

Before the theory of genetics was proposed, people selected outstanding plant and animal candidates based on their empirical experience of observing phenotypes. However, the phenotypes of traits were the results of the combined influence of genetic and environmental factors, therefore, genetic progress in plant and animal breeding was extremely slow due to the unquantified environmental contributions to phenotypic observations. In the mid-20th century, the BLUP (best linear unbiased prediction) model (Henderson 1975), which estimated the breeding values of each individual using the information from phenotypic observations, environmental records, and a relationship matrix derived from pedigree, was proposed and adopted in livestock breeding, and it has made great achievements in genetic improvements for agricultural economic traits. Nevertheless, the elements used to measure the genetic distance among individuals in the pedigree-based relationship matrix are values in theoretical expectations, could not capture the Mendelian sampling error fully, resulting in the same predictive performance for the full sibling individuals. With the development of sequencing technology, high density genetic markers across entire genome could be obtained, the genomic prediction, also known as genomic selection, was proposed subsequently by Meuwissen, Hayes, and Goddard (2001). Compared with traditional BLUP, the genetic markers could capture a greater proportion of Mendelian sampling error, thus genomic prediction is far more powerful at predicting almost all of the agricultural traits. However, how to model the enumerable phenotypic records with tens of thousands of genetic markers effectively to get a higher prediction performance has always been of prime concern in the domain of genomic prediction.

The most efficient strategy is to construct a relationship matrix using all the available markers, then replace the pedigree-based relationship matrix in the BLUP model, known as genomic BLUP (GBLUP), to obtain the genomic estimated breeding values (GEBVs), it is sample, robust and has been achieved in numerous software, e.g., **BLUPF90** (Misztal, Tsuruta, Strabel, Auvray, Druet, Lee et al. 2002), **DMU** (Madsen, Sørensen, Su, Damgaard, Thomsen, Labouriau et al. 2006). GBLUP model assumes that all markers have equal contributions to the phenotype. Obviously, this rough assumption is not quite appropriate for some of traits, especially for those that are controlled by several major genes, therefore the prediction accuracy usually varies across traits with different genetic architectures. Another more reasonable strategy is to fit Bayesian multiple regression model, known as individual level Bayesian model, which fits all the markers jointly and can flexibly assign different markers with various contributions that range from zero to infinity. Therefore the prior assumption would be more consistent with the practical genetic architecture of traits, which leads a high prediction performance compared with the GBLUP model. The big challenge with Bayesian regression model is the prior assumption on the contributions to the phenotype for each of the markers, because the prediction accuracy primarily depends on how close the prior assumption and the practical genetic architecture of the trait are. Statistical genetics researchers have constantly been devoted to optimizing the prior assumption, and a series of Bayesian methods have been proposed, collectively being called “BayesianAlphabet”. However, none of these methods consistently outperforms the others across different traits, because the genetic architecture is far more complex than the fitting capacity of a model (Wray, Wijmenga, Sullivan, Yang, and Visscher 2018). The package **BGLR** (Pérez and de los Campos 2014) is the most widely used tool to fit an individual level Bayesian model, but even so, the achieved methods in **BGLR** are limited, and it needs to be introduced with more advanced methods proposed lately.

In practice, it is always difficult to genotype all phenotypic individuals for a very large population, the individuals with effective phenotype observations, but without genotype information, are useless for genomic evaluation. Fernando, Dekkers, and Garrick (2014) proposed an advanced strategy for the first time, known as the single-step Bayesian regression model, which could utilize pedigree, genotype, and phenotype data simultaneously in a Bayesian linear model. The core idea of the single-step Bayesian regression model is to impute the genotype of the non-genotyped individuals in pedigree on the condition of the genotyped individuals by using the pedigree-based additive relationship matrix. As more phenotypic observations are used to fit the model, the prediction accuracy for genotyped individuals is higher than individual level Bayesian model, and for non-genotyped individuals, the prediction accuracy increases greatly compared with the pedigree-based BLUP model owing to the inclusion of the imputed genotype. Nevertheless, the problem is that there are only a few tools available to fit single-step Bayesian model, the package **JWAS** developed by Cheng, Fernando, and Garrick (2018), written in Julia, is the only one choice currently. However, the number of users and developers worldwide who use Julia is not comparable with the number who work in R.

To fit Bayesian regression model, the individual level data, including genome-wide genotype and one or several phenotypes measured on the same individuals, are required to be provided necessarily. However, the individual level data are sometimes not accessible to public due to some reasons of protection of personal privacy and legal or non-legal policies, especially in the area of human-related researches. Therefore, there are now continuously increasing GWAS summary statistical data publicly available on hundreds of complex traits, each of which consists of estimated effect sizes and sampling variance at millions of markers. The restricted access to individual level data has motivated statistical geneticists to develop new methodological frameworks that only require publicly available summary level data. At the last several years, Zhu and Stephens (2016) firstly introduced Bayesian regression into summary statistical analyses, and then Lloyd-Jones, Zeng, Sidorenko, Yengo, Moser, Kemper, Wang, Zheng, Magi, Esko, Metspalu, Wray, Goddard, Yang, and Visscher (2019) proposed a new advanced method, named as “SBayesR”, the results showed that SBayesR outperformed any of other existing non-Bayesian models in terms of prediction accuracy and with less cost for computing time. The summary data-based Bayesian model successfully transforms the determinant of computational complexity from the number of individuals into the number of markers, making it very promising for handing the data with large number of individuals that are genotyped by very few markers, for example, using chip arrays to genotype animals in the livestock breeding. As the summary level Bayesian model is still fresh to the public, the only one tool to fit summary level Bayesian model is **GCTB**, written in C++ by Zeng, de Vlaming, Wu, Robinson, Lloyd-Jones, Yengo, Yap, Xue, Sidorenko, McRae, Powell, Montgomery, Metspalu, Esko, Gibson, Wray, Visscher, and Yang (2018).

In addition to genomic prediction, Bayesian regression models can also be applied to genome-wide association studies (GWAS). Since it was firstly published in 2002 (Ozaki, Ohnishi, Iida, Sekine, Yamada, Tsunoda, Sato, Sato, Hori, Nakamura, and Tanaka 2002), GWAS has made great success in locating causal or candidate genes for human disease, as well as plant and animal agricultural economic traits, bringing new insights into understanding of the genetic architecture of various complex traits. Meantime, a series of advanced model have been developed for GWAS to overcome the population confounding problem that can cause the false positive associations. Some of these models include the general linear model in **Plink** (Purcell, Neale, Todd-Brown, Thomas, Ferreira, Bender, Maller, Sklar, de Bakker, Daly, and Sham 2007), the mixed linear model in **GCTA** (Yang, Lee, Goddard, and Visscher 2011) and **GEMMA** (Zhou and Stephens 2012), the multiple locus model “FarmCPU” in **rMVP** (Yin, Zhang, Tang, Xu, Yin, Zhang, Yuan, Zhu, Zhao, Li, and Liu 2021b). Although the statistical power to detect the true associations has increased consistently, all the models above are based on testing one marker, or only a small parts of markers jointly at a time, the significant associations explain only a small fraction of the genetic variance of quantitative traits, in addition, as the number of tests for those models can be very large, controlling the GWER (genome-wise error rate) results in very low power. In contrast, Bayesian regression models simultaneously fit all markers jointly as random effects, are able to account for most of the genetic variance, an advantage of this approach is that the power of detecting associations is not inversely related to the number of markers tested (Fernando, Toosi, Wolc, Garrick, and Dekkers 2017), thus, Bayesian regression models is an alternative option for GWAS analysis, but the developed tools are still not readily available.

As discussed above, although plenty of tools have been developed for different types of Bayesian regression models, they are written into various programming languages, the available models and methods are usually limited, and are not new enough for users or academic researchers. Moreover, the input file format and output results are quite different, making it expensive for comfortable usage in terms of effort and time expended. In this paper, we present **hibayes** (Yin, Zhang, Li, Zhao, and Liu 2021a), a comprehensive and user-friendly package developed on an open-source platform R (R Core Team 2013). The package contains the richest methods achieved thus far, and it is the only tool that can simultaneously fit three types of Bayesian models using individual level, summary level, and individual plus pedigree level (single-step) data for both genomic prediction and genome-wide association studies. It was designed to estimate joint effects and genetic parameters for a complex trait, including: (1) fixed effects and coefficients of covariates; (2) environmental random effects, and its corresponding variance; (3) genetic variance; (4) residual variance; (5) heritability; (6) genomic estimated breeding values for both genotyped and non-genotyped individuals; (7) marker effect size; (8) phenotype/genetic variance explained (PVE) by single or multiple markers; (9) posterior probability of association of the genomic window (WPPA); and (10) posterior inclusive probability (PIP). The functionalities are not limited, and we will keep on going in enriching **hibayes** with more features. **hibayes** is free software, licensed under the Apache License 2.0, openly available from the Comprehensive R Archive Network (CRAN) at https://CRAN.R-project.org/package=hibayes, and the latest version in development could be installed from GitHub at https://github.com/YinLiLin/hibayes.

## 2. Models and methods descriptions

In Bayesian statistics, inferences of unknown parameters of a model are based on their posterior distributions. Let ***θ*** denote all the unknown parameters in the model, then the full conditional posterior can be expressed by invoking Bayes’ law as:

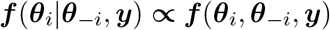

the joint density for the equation above can be written as

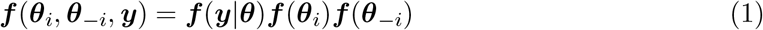

where ***f*** (***y***|***θ***) is the density function of the conditional distribution of ***y*** given the values of the unknowns specified by ***θ. f*** (***θ***_*i*_) and ***f*** (***θ***_*−i*_) are the densities of the prior distributions of ***θ***_*i*_ and ***θ***_*−i*_. By using MCMC techniques, all the elements in ***θ*** can be inferred from several number of iterations.

### 2.1. Individual level Bayesian model

Individual level Bayesian model is essentially a version of multiple linear regression model, which describes the phenotypic observations as a function of hundreds of thousands of variables, including fixed effects, covariates, environmental random effects, and high dimension of genetic markers. The model could be mathematically formulated as:

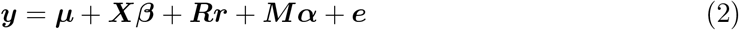

where ***y*** is vector of the phenotypic observations, ***X*** represents the designed model matrix for fixed effects and covariates, ***R*** is the index matrix for the environmental random effects, ***M*** is the genotype covariate matrix with a dimension of ***m*** * ***n***, where ***m*** and ***n*** are the number of individuals and genetic markers, respectively, and ***e*** is the vector of residuals. Let 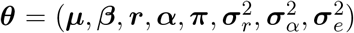 denote all the unknown hyper-parameters in the model of Equation 2, including intercept, coefficients of fixed effects and covariates, the environmental effects, the marker effects, the mixing proportions of the mixture of normal terms, the variance of environmental effects, the variance of marker effects, and the residual variance. As shown in the Equation 1, the posterior distribution of one parameter is on the condition of the other parameters, thus we first need to assign all the unknown parameters with prior values at the beginning to derive the full conditional distributions. Let

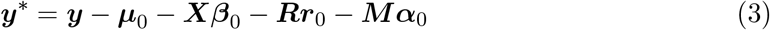

where ***µ***_0_, ***β***_0_, ***r***_0_, ***α***_0_ are the initialized start values for the unknown model parameters ***µ, β, r, α***, respectively. In package **hibayes**, the intercept ***µ***_0_ is assigned to be the average of dependent variable ***y***, and ***β***_0_, ***r***_0_, and ***α***_0_ are initialized in zeros at the start of the MCMC iteration.

#### Fixed effects and covariates

For fixed effects, we can not fit it in the model directly, it should be converted into a designed matrix in advance, then combined with covariates by column into the final model matrix to fit model. In **hibayes**, each column of the model matrix is treated as an independent random variable, and follows a normal distribution, the full conditional distribution can be formulated on condition of other parameters as follows:

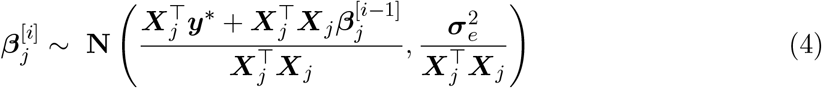

where 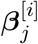 is the sampled value of ***j***_*th*_ variable for the ***i***_*th*_ iteration, and 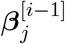 is for the previous iteration. ***X***_*j*_ represents the ***j***_*th*_ column of the model matrix ***X***. The corresponding residuals for 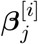 are

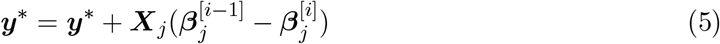

repeat Equation 4 and 5 to update the posterior estimations of ***β*** for the current iteration.

#### Environmental random effects

When an environmental factor needs to be considered in the model, the levels of the records for this factor can not cover all the possibilities of the entire population, then it is generally treated as an random term in the model, which follows a normal distribution 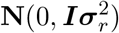. The conditional solution based on the BLUP frame for the effects ***r*** of environmental random term is

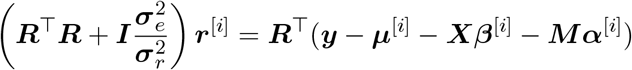

where the superscript [*i*] represents the *i*_*th*_ iteration of the MCMC process. To obtain the right side of the equation above is usually time-expensive for a large number of iterations, however, it can be derived from the estimations of previous iteration as follows:

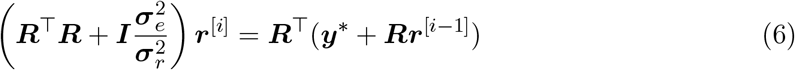

where the superscript [*i −* 1] represents the estimations of previous iteration. To get the posterior estimations ***r***^[*i*]^, two types of sampler algorithms can be taken into consideration, the first is the single-site Gibbs sampler (Sorensen and Gianola 2002), which draws parameters from each conditional distribution with density ***p***(***r***_*k*_|***r***_*−k*_), ***k*** = **1**, …, ***n***_*r*_, where ***n***_*r*_ is the number of elements in the environmental random effect ***r***. By simplifying the Equation 6 to ***Cr***^[*i*]^ = ***B***, then we can formulate single-site Gibbs sampler strategy as follows:

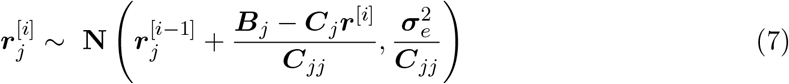

where 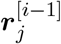 represents the ***j***_*th*_ element of ***r*** at the MCMC iteration *i −* 1, ***B***_*j*_ is the ***j***_*th*_ element of ***B***, and ***C***_*jj*_ and ***C***_*j*_ are the ***j***_*th*_ diagonal element and the column ***j*** of matrix ***C***, respectively. Obviously it can not be processed in parallel, but the big advantage is that there is no need to compute the inverse of matrix ***C***; the second algorithm is the block Gibbs sampler (García-Cortés and Sorensen 1996; Lund and Jensen 1999), which draws unknown parameters jointly from the a multivariate distribution, and it can be written as

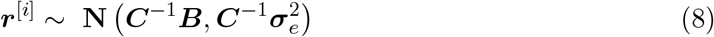

this requires the inverse of matrix ***C***, making it to be competent in handing sparse matrix. For the Equation 6, the number of levels for the environmental random effect is usually small, and the left-hand side of the equation is very sparse, therefore, both single-site Gibbs sampler and block Gibbs sampler are applicable. After the environmental random effects are sampled successfully, then the corresponding residuals of model could be updated by

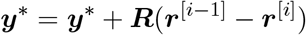

The variance of environmental random effects 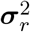 is assumed to follow a scaled-inverse chisquare distribution expressed as follows:

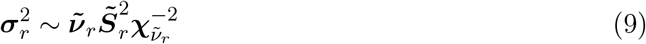

where 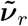 and 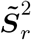 are the degree of freedom and the scale factor of the inverse chi-square distribution, respectively, which can be derived from

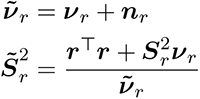

where ***ν***_*r*_ = 4 by default, 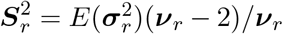, and 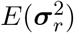 is initialized to be the variance 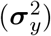 of dependent variable ***y***.

#### Genetic marker effects

Similar to Equation 4, we can also draw the posterior effect size of genetic marker from a normal distribution on condition of the sampled value of all other variables and the data, but differently, the effect size of markers are allocated into different groups, each group takes different categorical distribution, and the expected value of the marker is weighted by a combination of its corresponding group variance and residual variance, it can be mathematically formulated as follows:

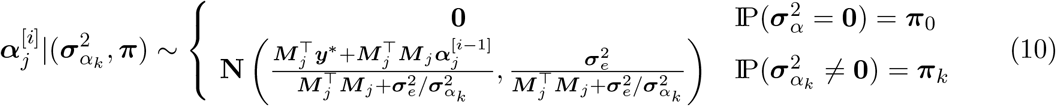

where where 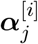 represents the ***j***_*th*_ element of the marker effect size ***α*** at ***i***_*th*_ MCMC iteration, ***π***_0_ and ***π***_*k*_ are the sub-elements of ***π, π***_0_ is the probability of stepping into zero effect size for ***j***_*th*_ marker (the proportion of markers in zero effect size), ***π***_*k*_ is the probability of stepping into the ***k***_*th*_ categorical distribution, of which the variance is 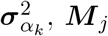 represents the coded genotype vector for ***j***_*th*_ markers (column ***j*** of matrix ***M***), and IP is the probability. The variance 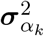 reflects the contribution of ***j***_*th*_ marker to the dependent variable ***y***, thus the assumption of the distribution of 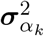 for all markers is related closely to the complexity of the genetic architecture of a trait. How to give a more appropriate assumption of the distribution of 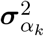 has long been a hot and difficult topic in the domain of genomic prediction during recent decades, therefore, a series of Bayesian methods has been proposed aimed at improving the prediction accuracy, collectively being called the “BayesianAlphabet”. However, more and more evidences has shown that none of any method can always outperform the others across different traits (Yin, Zhang, Zhou, Yuan, Zhao, Li, and Liu 2020). Here in **hibayes**, we have achieved some typical methods, including BayesRR, BayesA, BayesB(pi), BayesC(pi), BayesLASSO, and some advanced methods used most frequently, including BSLMM and BayesR. The rough descriptions for the assumption on the distribution of 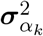 for the methods mentioned above are in Table 1 and as follows:

**Table 1:** Assumption of effect size distribution of markers for the methods in **hibayes**.

| Model | Formula | Joint distribution | Reference |
| --- | --- | --- | --- |
| BayesRR | $\alpha_j \sim \mathbf{N}(0, \sigma_\alpha^2)$ | normal | de los Campos, Hickey, Pong-Wong, Daetwyler, and Calus (2013) |
| BayesA | $\alpha_j \sim \mathbf{N}(0, \sigma_{\alpha_j}^2)$<br>$\sigma_{\alpha_j}^2 \sim \chi^{-2}(\nu, S)$ | t | Meuwissen *et al.* (2001) |
| BayesB | $\alpha_j \sim 0.05 \mathbf{N}(0, \sigma_{\alpha_j}^2) + 0.95\delta_0$<br>$\sigma_{\alpha_j}^2 \sim \chi^{-2}(\nu, S)$ | point-t | Meuwissen *et al.* (2001) |
| BayesBpi | $\alpha_j \sim (1 - \pi) \mathbf{N}(0, \sigma_{\alpha_j}^2) + \pi\delta_0$<br>$\sigma_{\alpha_j}^2 \sim \chi^{-2}(\nu, S)$ | point-t | Meuwissen *et al.* (2001) |
| BayesC | $\alpha_j \sim 0.05 \mathbf{N}(0, \sigma_\alpha^2) + 0.95\delta_0$ | t mixture | Habier, Fernando, Kizilkaya, and Garrick (2011) |
| BayesCpi | $\alpha_j \sim (1 - \pi) \mathbf{N}(0, \sigma_\alpha^2) + \pi\delta_0$ | t mixture | Habier *et al.* (2011) |
| BayesL | $\alpha_j \sim \mathbf{N}(0, \sigma_{\alpha_j}^2)$<br>$\sigma_{\alpha_j}^2 \sim \text{Expon}(\lambda^2/2)$ | double exponential | Yi and Xu (2008) |
| BSLMM | $\alpha_j \sim (1 - \pi) \mathbf{N}(0, \sigma_{\alpha_1}^2) + \pi \mathbf{N}(0, \sigma_{\alpha_1}^2 + \sigma_{\alpha_2}^2)$ | normal mixture | Zhou, Carbonetto, and Stephens (2013) |
| BayesR | $\alpha_j \sim \pi_1 \mathbf{N}(0, 10^{-2}\sigma_\alpha^2) + \pi_2 \mathbf{N}(0, 10^{-3}\sigma_\alpha^2) + \pi_3 \mathbf{N}(0, 10^{-4}\sigma_\alpha^2) + (1 - \pi_1 - \pi_2 - \pi_3)\delta_0$ | point-normal mixture | Moser, Lee, Hayes, Goddard, Wray, and Visscher (2015) |

Note: $\delta_0$ represents the effect size equals to zero, $\nu$ and $S$ are the degree freedom and scale parameter for inverse chi-square distribution, $t$ represents student's t-distribution.

**BayesRR:** Bayesian Ridge Regression, all markers have non-zero effects and share the same variance, equal to RRBLUP (ridge regression BLUP) or GBLUP (genomic BLUP).

**BayesA:** all markers have non-zero effects, and take different variances which follow an inverse chi-square distribution.

**BayesB:** most of the markers have zero effects (***π***_0_, ***π***_0_ = 0.95 by default), only a small proportion of markers (1 *−* ***π***_0_) have non-zero effects, and take different variances which follow an inverse chi-square distribution.

**BayesBpi:** the same as **BayesB**, but ***π***_0_ is not fixed, will be estimated in the iterative process.

**BayesC:** most of the markers have zero effects (***π***_0_, ***π***_0_ = 0.95 by default), only a small proportion of markers (1 *−* ***π***_0_) have non-zero effects, and share the same variance.

**BayesCpi:** the same as **BayesC**, but ***π***_0_ is not fixed, will be estimated in the iterative process.

**BayesL:** BayesLASSO, all markers have non-zero effects, and take different variances which follow an exponential distribution.

**BSLMM:** all markers have non-zero effects, and take the same variance, which can be captured by GBLUP model, but a small proportion of markers has an additional shared variance.

**BayesR:** only a small proportion of markers have non-zero effects, and the markers are allocated into different groups of normal distributions, and the relative variance for each normal distribution is fixed.

Before sampling the posterior effect size for the ***j***_***th***_ genetic marker by Equation 10, it must be known to which categorical distribution should the ***j***_***th***_ genetic marker belongs. Thus, we first need to calculate all likelihoods assuming the considered genetic marker ***j*** being in 1 of the ***n***_*π*_ (the number of elements of ***π***) normal distributions at a time with the respective probability ***π***. The likelihood that the ***j***_***th***_ genetic marker is in distribution ***k*** is

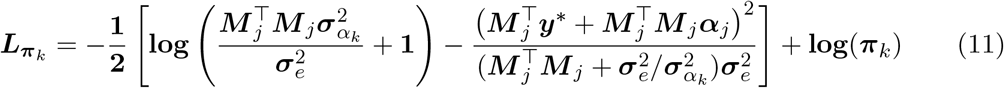

where ***π***_*k*_ is the probability for ***k***_***th***_ categorical distribution (or the ***k***_***th***_ value in the vector of ***π***). The detailed derivation for 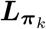 can be found in the paper published by Lloyd-Jones *et al*. (2019).

Then, as described by Erbe, Hayes, Matukumalli, Goswami, Bowman, Reich, Mason, and Goddard (2012), the probability that marker ***j*** is in distribution ***k*** is

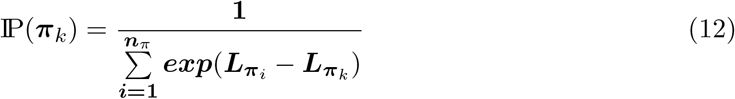

Based on these probabilities, we can select the normal distribution to draw the marker effect from using a uniform random variate *U* (0, 1), using the probabilities of the marker being in each of the categorical distributions. Once the distribution for marker ***j*** has been confirmed, the effect size can be sampled by Equation 10. Then update the residuals as follows:

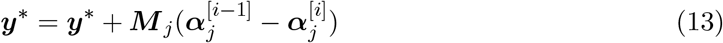

By repeating the Equation 11 → 12 → 10 → 13, the effect size for all markers can be updated, and we can record the number of markers allocated into different categorical distributions for the current iteration:

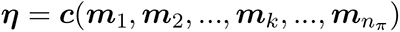

Where 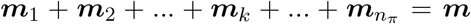, and ***m*** is the total number of markers. If ***π*** needs to be estimated, the posterior ***π*** can be sampled from a *Beta* distribution, taking 2 categorical distributions for an example, let 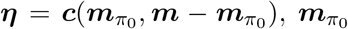 is the number of genetic markers in zero effect size, then we have

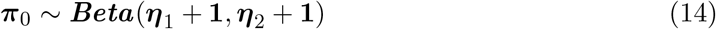

Or it can also be sampled from a *Dirichlet* distribution

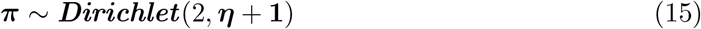

It should be noted that if the number of categorical distributions is bigger than 2 (***n***_*π*_ > 2), only the *Dirichlet* distribution can be used for sampling, for example, BayesR.

The variances of effect size of all markers are assumed to follow an inverse chi-square distribution as the Equation 9, which can be sampled on condition of the updated marker effect ***α***.

For the methods that each marker has unique variance, e.g. BayesA, BayesB(pi), BayesLASSO, the sampling procedure for variance should be implemented independently following sampling marker effect, and the equation is

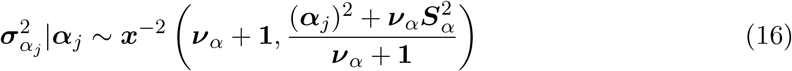

where the degree of freedom ***ν***_*α*_ = **4**, and the scale parameter 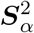 is

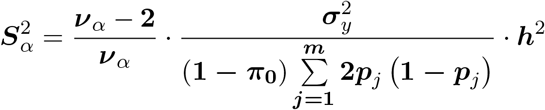

where ***h***^2^ is the initialized heritability of the trait, which is set to be 0.5 in **hibayes, *p***_***k***_ is the frequency of allele at ***j***_*th*_ genetic marker.

The sample strategies for different methods are slightly different. For the methods that markers are allocated into different categorical distributions, e.g., BayesRR, BayesC(pi), the sampling procedure for variance is implemented at the end of sampling effect of the last genetic marker, and it is not necessary to implement sampling for every single marker, but for different groups of categories, except for the group in zero effect size, the distribution can be referred as

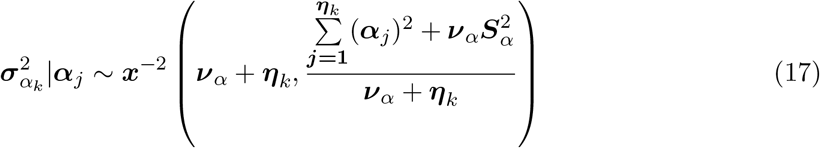

where ***η***_*k*_ is the number of markers allocated into the ***k***_*th*_ categorical distribution.

It should be pointed out that, for BayesR method, the variances for the categorical distributions are assigned on the scale of 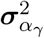, the scale ***γ*** = ***c***(0.0001, 0.001, 0.01), thus we do not need to draw the variance for different categorical distributions, but only sample the variance 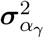 once as follows:

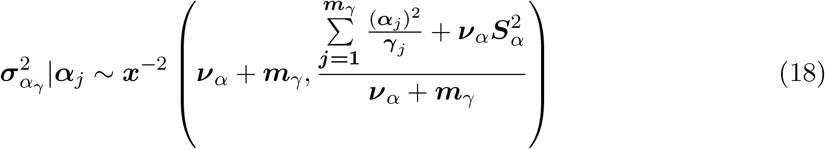

Where 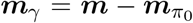, and ***γ***_*j*_ is the scaled value of the categorical distribution for the ***j***_*th*_ genetic marker. In the Equation 10 and 11, 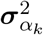 should be replaced as 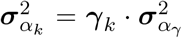 for BayesR method.

For BSLMM method, as described by Zhou *et al*. (2013), the linear model equation has an additional term compared with Equation 2:

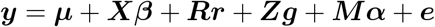

where ***Z*** is the designed matrix, ***g*** is a vector of random effects, also known as GEBV (genomic estimated breeding value), referring to standard terminology from GBLUP model, it is assumed to follow a normal distribution 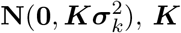 is the additive genomic relationship matrix (GRM) that is derived from genotype information of all individuals, ***α*** is the vector of additional genetic effects that can not be captured by ***g*** for a small proportion of genetic markers, which come from one normal distribution as the same with BayesCpi. As described by VanRaden (2008), the GRM could be constructed using the following equation:

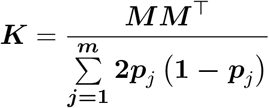

where ***m*** is the number of genetic markers, ***p***_***k***_ is the frequency of allele ***A***_**1**_ at ***j***_*th*_ genetic marker, ***M*** is the additive marker covariate matrix with elements of **2 *−* 2*p***_*j*_, **1 *−* 2*p***_*j*_, and ***−*2*p***_*j*_ for ***A***_**1**_***A***_**1**_, ***A***_**1**_***A***_**2**_, and ***A***_**2**_***A***_**2**_, respectively. In **hibayes** package, we do not strictly follow the sampling strategy proposed by Zhou *et al*. (2013), we implement a procedure on the combination of single-site Gibbs sampler (Equation 7) and BayesCpi sampler to achieve the model assumption of BSLMM, this means that the BSLMM method is an extension of BayesCpi with an additional multivariate normal random term ***g***. Similar to the Equation 6, the solution to ***g*** is

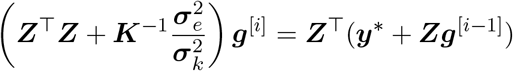

to construct the equation above, we need to compute the inverse of ***K*** matrix. However, ***K*** is not always invertible for some reasons (e.g., not positive defined), so we have achieved three types of algorithms to address this problem in package **hibayes**: Cholesky decomposition, lower–upper (LU) decomposition, and ridge regression. As ***K***^*−*1^ is no longer sparse, the block Gibbs sampler (García-Cortés and Sorensen 1996; Lund and Jensen 1999), which requires the inverse of the left-hand side of the equation for every single iteration, would be time-consuming within the MCMC process, therefore we use the single-site Gibbs sampler (Sorensen and Gianola 2002) to obtain the solution of ***g*** in package **hibayes**. 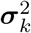 follows a scaled inverse chi-square distribution as Equation 9, with degree of freedom and scale parameter:

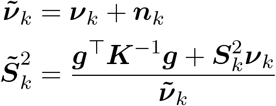

where ***n***_*k*_ is the number of elements in ***g, ν***_*k*_ = 4, and

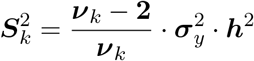

different with BayesCpi, the final marker effect size of BSLMM method includes two parts:

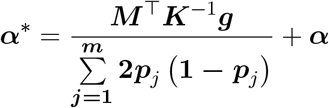

the first part transforms the GEBVs into marker effects following the paper published by Aliloo, Pryce, González-Recio, Cocks, Goddard, and Hayes (2017). The operation of multiplication for two big matrices in the equation above is extremely time-consuming, however, this step is implemented only once at the end of the MCMC iteration and, therefore, would not be a big problem.

#### Residual effects

The residuals of the linear model will be updated subsequently once any of model parameter (***µ, β, r, α***) is sampled from its corresponding full conditional distribution. The variance of residuals 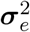 follows the same distribution as Equation 9, the degree of freedom and scale parameter are as follows:

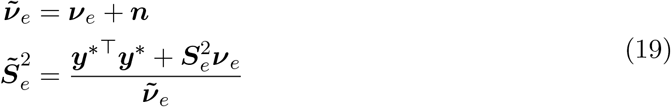

where ***ν***_*e*_ = *−*2, and 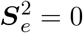 by default in **hibayes**.

### 2.2. Summary level Bayesian model

To fit individual level model, the individual level data, including genome-wide genotype and one or multiple phenotypes measured on the same individuals, should be provided. However, the individual level data are sometimes not accessible to public due to personal privacy, legal or non-legal policies, especially when related to research on humans. In addition to individual level data, there are now continuously increasing GWAS summary statistics available on hundreds of complex traits, each consisting of estimated effect sizes and sampling variance at millions of markers. Restricted access to individual level data has motivated methodological frameworks that only require publicly available summary level data, one of which is the Bayesian multiple regression summary statistics proposed in recent years (Zhu and Stephens 2016; Lloyd-Jones *et al*. 2019). The summary level Bayesian model is inferred from individual level model, so let

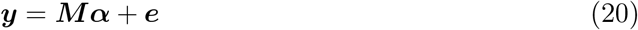

different with Equation 2, here ***y*** is the prior adjusted phenotype by the fixed effects, covariates, environmental random effects, rather than the original phenotype, the reason for the adjustment is that the public summary data only includes summary information for genetic markers, and we can not directly access the recorded environmental factors, thus the unknown parameters of summary level Bayesian model 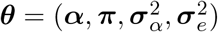.

Let 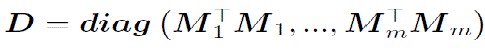, by multiplying Equation 20 by ***D***^*−*1^***M*** ^*T*^ to arrive at

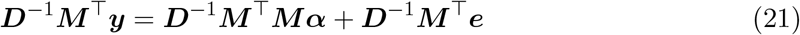

noting that the variance-covariance matrix (***V***) of all genetic markers could be written as

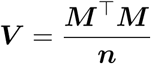

***V*** could be calculated from the publicly available reference genotype panel. Then we can rewrite the Equation 21 as

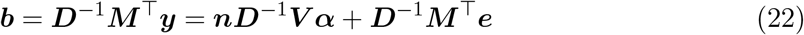

where ***b*** is vector of the marginal effects (also known as regression coefficients), which can be obtained from summary data directly. Therefore, we can successfully transform the individual level Bayesian model into the summary level Bayesian model. Noting that if the ***V*** matrix is derived from the same genotype panel with summary data, then the estimated results of unknown parameters of summary level Bayesian model will be the same as individual level Bayesian model in theory.

As described in Section 2.1, to sample the marker effect size ***α***_*j*_, we need to know to which categorical distribution it belongs, thus the first step is to calculate likelihoods for all categorical distributions for marker ***j***. Referring to Equation 11, it requires the components: 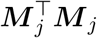, and the prior values of 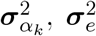, and 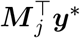. 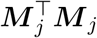 is the diagonal element of ***M*** ^*T*^***M***, and it can be calculated as

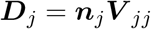

where ***V*** _*jj*_ is the ***j***_*th*_ diagonal element of ***V***, ***n***_*j*_ is the effective number of individuals for marker ***j***, which can be obtained directly from summary data, and as we know that if the marker is in Hardy-Weinberg equilibrium, then we can also derive ***D***_*j*_ as follows:

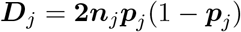

where ***p***_*j*_ is the allele frequency for marker ***j***, and can be calculated from the reference genotype panel.

The prior values of 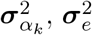 can be obtained from scaling the phenotype variance 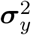, and 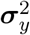 can be derived from summary data as follows:

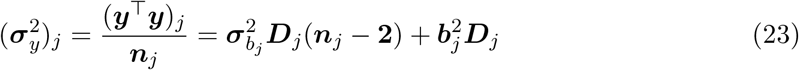

where 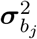 is the square of the standard error of ***b***_*j*_, and it can also be obtained directly from summary data, then taking the median over the set of 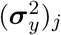 can get reliable estimation of 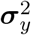 (Yang,, Ferreira, Morris, Medland, Madden, Heath, Martin, Montgomery, Weedon, Loos, Frayling, McCarthy, Hirschhorn, Goddard, and and 2012). The main challenge for summary level Bayesian model is to get 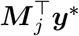, where ***y***^*^ is the vector of residuals on condition of other model parameters, however, the phenotype observations ***y*** is not available in summary data,thus ***y***^*^ can not be accessed and updated in MCMC iterations, let

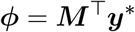

then 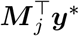 is the ***j***_*th*_ element of ***ϕ***, which is ***ϕ***_*j*_. At the beginning of the MCMC iteration,***α*** is initialized in a vector of zeros, ***y***^*^ = ***y***, then the start values for ***ϕ*** can be expressed as

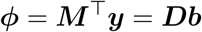

now we can calculate the likelihoods, and compute the probabilities for all categorical distributions following the Equation 12, then follow the steps described in Section 2.1 to draw the posterior marker effect size ***α***_*j*_.

Once ***α***_*j*_ is updated, the next step is to update the residuals ***y***^*^ following the Equation 13, as ***y***^*^ can not be accessed, we multiply Equation 13 by ***M*** ^*T*^:

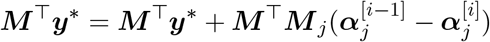

which can be rewritten as

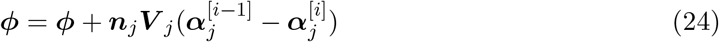

where ***V*** _*j*_ is a vector of the ***j***_*th*_ column of ***V***. By the transformation above, we just repeatedly compute ***ϕ*** for each update of the marker effect size to keep the MCMC iteration going correctly, and those processes successfully cut off the dependency on ***y***^*^.

The sampler strategies for ***π*** and 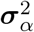 are the same with individual level Bayesian model referring to Equation 14, 15 and Equation 16, 17, 18. The other two parameters of importance are 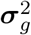 and 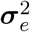, these values are used to estimate heritability ***h***^2^. 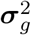 is the genetic variance, which is defined as the variance of GEBVs, then we have

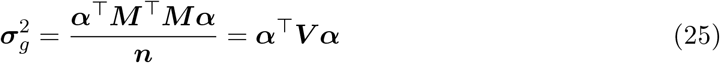

the operation of multiplication for a large matrix is expensive within the MCMC process. From Equation 20, we have ***Mα*** = ***y*** *−* ***y***^*^, multiplied by ***M*** ^*T*^, then

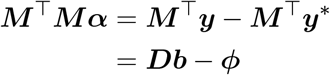

we can rewrite

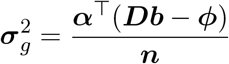

now it becomes the multiplication of two vectors, which will be much more efficient than Equation 25.

For the residual variance 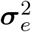, similarly, we have

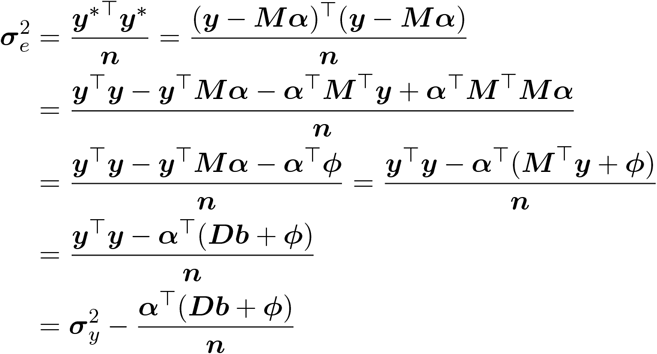

where 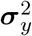 can be computed by Equation 23. The final global 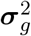 and 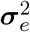 can be sampled from the scaled-inverse chi-square distribution by integrating equations above into Equation 9.

The available methods for summary level Bayesian model are nearly the same as for individual level Bayesian model, except for BSLMM, because the individual genotype is inaccessible to construct GRM. Additionally, we have implemented a conjugate gradient (CG) algorithm in **hibayes** to solve the Equation 22 to obtain the marker effect size directly without running MCMC process, it equals to SBLUP (summary level BLUP) in theory (Robinson, Kleinman, Graff, Vinkhuyzen, Couper, Miller, Peyrot, Abdellaoui, Zietsch, Nolte et al. 2017). The CG algorithm requires a ridge regression value ***λ***, which should be provided as ***λ*** = ***m***\*(**1***/****h***^2^ *−***1**), where ***h***^2^ is the heritability of the trait and can be estimated by LD score regression using **LDSC** software (Bulik-Sullivan, Loh, Finucane, Ripke, Yang, Patterson, Daly, Price, and Neale 2015).

Looking back at the relative theoretics of summary level Bayesian model, it is easy to find that how to fast compute and to store the variance-covariance matrix ***V*** is a big problem, the dimension of square matrix ***V*** equals to the number of genetic markers in analysis, although it is narrowly acceptable for chip array data at a level of several tens of thousands of markers, the high density markers from sequencing would be a big challenge and a bottleneck faced for summary level Bayesian model. However, typically this problem can be addressed by using a fixed 1–10Mb window approach, as in SBLUP (Robinson et al. 2017) or LDpred (Vilhjálmsson, Yang, Finucane, Gusev, Lindström, Ripke, Genovese, Loh, Bhatia, Do et al. 2015), which sets LD correlation values outside this window to zero, or using a shrunk LD matrix proposed by Zhu and Stephens (2017). In **hibayes** package, we use a chi-square threshold 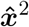 to make ***V*** matrix to be sparse, if the condition 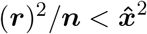 (***r*** is the Pearson correlation coefficient of two markers) is met, then the covariance for those two markers will be set to zero. Additionally, we have added an option in **hibayes** to let users choose whether to compute the covariance among chromosomes. Therefore, **hibayes** can compute totally 4 types of variance-covariance matrix: genome-wide full dense matrix, genome-wide sparse matrix, chromosome-wide full dense matrix, or chromosome-wide sparse matrix.

It should be noted that, if using a sparse matrix to fit summary level Bayesian model, some of elements of ***ϕ*** in Equation 24 are not updated, then the mean of posterior distribution in Equation 10 for the markers relevant to those elements would be biased, which may sometimes cause the MCMC process to “blow up” in certain situations, we recommend adjusting the chisquare threshold 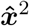 to address this problem.

### 2.3. Single-step Bayesian model

It is always difficult to genotype all individuals of a large population in practice, meaning that the individuals with effective phenotype observations but without genotype information are useless for genomic evaluation. Therefore, in 2010, a single-step BLUP model (SSBLUP), which can simultaneously integrate pedigree, genotype, and phenotype data, was proposed to maximize the utilization of available information, making it possible to connect all phenotypic observations for both genotyped and non-genotyped individuals (Christensen and Lund 2010; Aguilar, Misztal, Johnson, Legarra, Tsuruta, and Lawlor 2010). However, on the one hand, SSBLUP model requires GRM and its inverse of all genotype individuals, which is an inefficient process, because GRM is very dense and it grows in size as more individuals are genotyped. On the other hand, SSBLUP model assumes that all genetic markers have the equal contributions to the phenotype, which is inconsistent with the real genetic architecture of traits, especially for a trait controlled by several major genes. Subsequently, single-step Bayesian regression (SSBR) model was proposed by Fernando *et al*. (2014); Fernando, Cheng, Golden, and Garrick (2016), SSBR is an extension of individual level Bayesian model, and can simultaneously fit pedigree, genotype, and phenotype data in a Bayesian linear model for the first time, the mathematical formula for SSBR model is

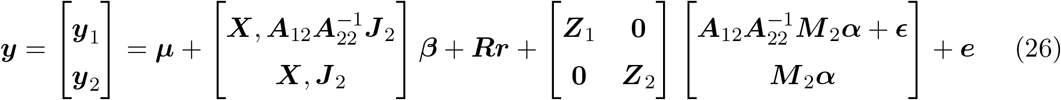

the unknown parameters 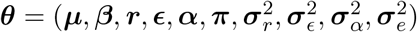, where ***y***_1_ and ***y***_2_ are the phenotypic records for non-genotyped and genotyped individuals, respectively, ***J*** _2_ = *−***1, *A***_12_ is the pedigree based additive relationship matrix between non-genotyped and genotyped individuals, 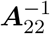 is the inverse of the pedigree based additive relationship matrix between genotyped individuals, ***M*** _2_ is the genotype covariate matrix for genotyped individuals, ***E*** is the vector of imputation residuals for non-genotyped individuals, and other symbols are the same as Equation 2.

The core idea of the SSBR model is to impute the genotype of the non-genotyped individuals in pedigree on condition of the genotyped individuals by using the pedigree based additive relationship matrix (***A***). However, to construct ***A***_12_ and to compute the inverse of ***A***_22_ in Equation 26 are not such efficient with the increasing size of pedigree. Fortunately, Fernando *et al*. (2014) proved that the imputed markers can be obtained efficiently, using partitioned inverse results, by solving the easily formed very sparse system:

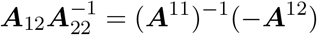

where ***A***^11^ is the partition of ***A***^*−*1^ for non-genotyped individuals, ***A***^12^ is the partition of ***A***^*−*1^between non-genotyped and genotyped individuals. As ***A***^11^ is very sparse, it will be quite fast to get the solution on left hand side by solving the sparse linear system on the right side, thus ***A*** is no longer required, only ***A***^*−*1^. In 1976, Henderson presented a simple procedure to derive ***A***^*−*1^ from pedigree directly without inverting ***A*** matrix (Henderson 1976). The inverse of ***A*** matrix can be written as:

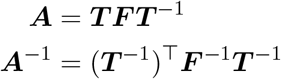

where ***T*** is a lower triangular matrix and ***F*** is a diagonal matrix. ***T*** ^*−*1^ is a lower triangular matrix with ones in the diagonal, and the only non-zero elements to the left of the diagonal in the row for the individuals are −0.5 for columns corresponding to the known parents. Regardless of the level of inbreeding, the diagonal elements of ***F*** ^*−*1^ are either 2 or 4/3 or 1, if both or one or no parents are known, respectively. The ***A***^*−*1^ matrix can be derived from pedigree using the following R codes:

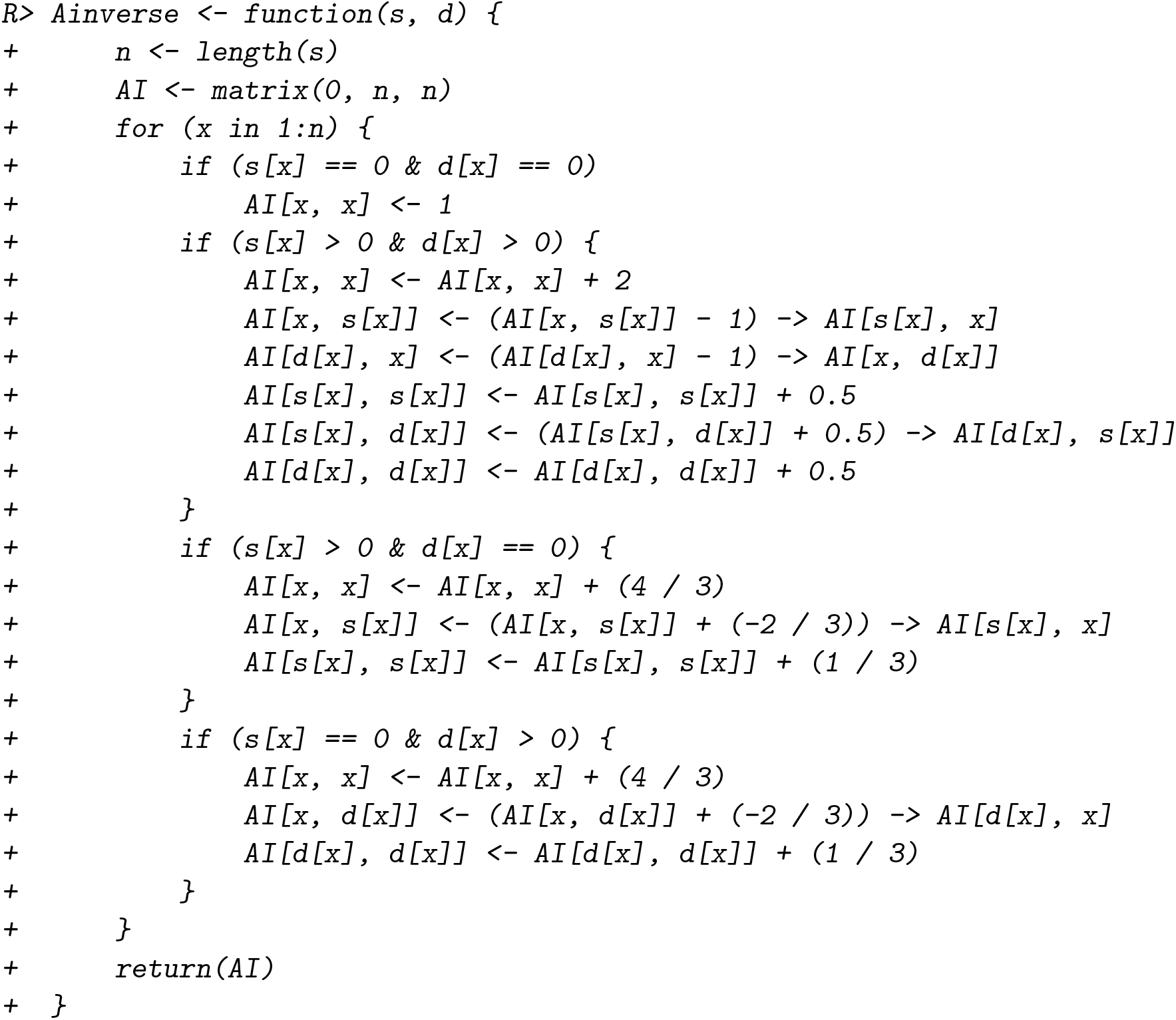

Once the genotype imputation is done successfully, it is easy to fit single-step Bayesian model, which is almost the same as individual level Bayesian model, except for the imputation residuals ***ϵ***, which is assumed to follow a multivariate normal distribution. Similar to Equation 6, the conditional distribution of ***ϵ*** on the sampled value of all other variables is given by the solution of the following system:

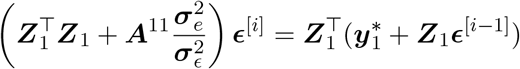

where 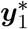 is the sub-vector of ***y***^*^ (residuals of the model) for non-genotyped individuals. As discussed in Equation 6, we can draw ***ϵ***^[*i*]^ by two types of algorithms, here ***A***^11^ is pretty sparse, it will be efficient to obtain the inverse of the left hand side of the equation above, thus block sampler is more appropriate for this situation. 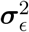 is the variance of imputation residuals ***ϵ***, the full-conditional posterior for 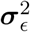 is a scaled inverse chi-square distribution as Equation 9, with degree of freedom and scale parameter:

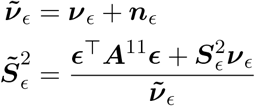

where ***n***_***ϵ***_ is the number of elements in ***E, ν***_***ϵ***_ = 4, and 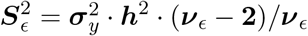.

Once all the values in vector of ***ϵ*** are sampled, then update the residuals 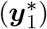 for non-genotyped individuals as follows:

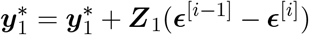

and keep the residuals 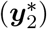 of genotyped individuals unchanged. The sampler strategy for other unknown parameters in ***θ*** is the same with individual level Bayesian model, and can be referred in Section 2.1.

The available methods for single-step Bayesian model in **hibayes** is nearly the same with individual level Bayesian model as described in Table 1, except for BSLMM, as the BSLMM method requires the GRM of all individuals, however, the imputed genotype for non-genotyped individuals can not be used directly to construct GRM, and to compute the marker effect size, because there are no specific imputation residuals for markers, but for individuals, thus the estimated GRM and marker effect size would be biased merely using the imputed genotype.

It should be pointed out that for SSBR model, the number of predicted individuals depends on the number of unique individuals in pedigree, if all the individuals with phenotypic records have been genotyped, then the imputation residuals can not be estimated, although the non-genotyped individuals in pedigree still could be predicted in **hibayes**, the prediction accuracy should be validated further with real data.

### 2.4. Genomic prediction and Genome-wide association studies

#### Genomic prediction

For genomic selection, the main purpose is to obtain the individual’s GEBV, which reflects the difference of a individual on genetic performance over the average level of the whole population. As discussed in the section above, Bayesian regression models only estimate the effect size of markers across the entire genome, in order to obtain the GEBVs, we need the individual level genotype. If we have the genotype in hand, then generally it’s easy to derive GEBVs from the following equation:

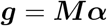

this can be accomplished by the function”`--score`” of **PLINK** using the prepared genotype file in binary format and marker effect size file in text format. However, for single-step Bayesian model, as described by Fernando *et al*. (2014), the GEBVs includes three parts as follows:

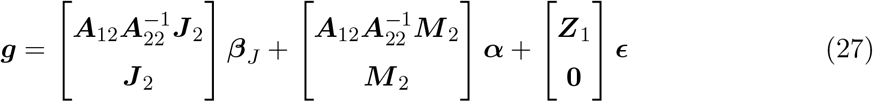

where the symbols can be referred to Equation 26, the first part comes from the estimated coefficient, the second part is derived from genotype, and the third part includes the imputation residuals for non-genotyped individuals. There is no software or pipeline that can be used directly to accomplish the above calculation, and general speaking, it is not that easy for users to implement this manually. Therefore, in **hibayes**, we have reported GEBVs directly for all individuals in the final returned list. For individual level Bayesian model, both phenotypic individuals or non-phenotypic (marked as ‘NA’) individuals that have genotype will be predicted. For single-step Bayesian model, all individuals in pedigree, including genotyped or non-genotyped individuals, will be predicted in the final step. But for summary level Bayesian model, as the individual level genotype is not compulsorily required as an input, so it should be accomplished manually by users.

#### Genome-wide association studies

Bayesian regression model can not only be used for genomic prediction, but also could be applied to genome-wide association studies to locate candidate genes of a trait (Yi, George, and Allison 2003; Fan, Onteru, Du, Garrick, Stalder, and Rothschild 2011). Given such a model where ***π*** is close to one, the posterior probability that ***α***_*j*_ is nonzero for at least one marker ***j*** in a window or segment can be used to make inferences on the presence of QTL (quantitative trait locus) in that segment. We refer to this probability as the window posterior probability of association (WPPA). The underlying assumption here is that if a genomic window contains a QTL, one or more markers in that window will have nonzero ***α***_*j*_. Thus, WPPA, which is estimated by counting the number of MCMC samples in which ***α***_*j*_ is nonzero for at least one marker ***j*** in the window, can be used as a proxy for the posterior probability that the genomic window contains a QTL. WPPA can be formulated as

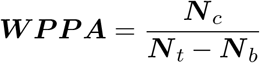

where ***N*** _*t*_ is the total number of iteration of MCMC process, ***N*** _*b*_ is the number of discarded iteration at front of MCMC process, and ***N*** _*c*_ is the counted number of times that ***α***_*j*_ is nonzero for at least one marker ***α***_*j*_ in the window after the discarded iterations.

Also, **hibayes** provides the posterior probability that ***α***_*j*_ is nonzero for each individual markers, which is known as posterior inclusive probability (PIP), it is the same with WPPA when the window size equals to 1, and can be used as a complementary result for more detailed location of causal markers.

## 3. The R package hibayes

As R has become the one of the most widely used languages for statistical computing and graphics, with a large number of users worldwide, we developed **hibayes** on the R platform. However, as it is described about the details of Bayesian regression model in the above section, there are huge number of iterations for parameter estimation, obviously it is not a good decision to write Bayesian regression model into pure R language. Therefore, we accomplished the core parts which take over the most of computation time of Bayesian regression model into C++ language by the aid of the packages **Rcpp** (Eddelbuettel and François 2011) and **RcppArmadillo** (Eddelbuettel and Sanderson 2014), all parallelizable parts were sped up by **OpenMP** (Dagum and Menon 1998), some basic vector operations were enhanced by calling corresponding functions in **LAPACK** (Anderson, Bai, Bischof, Blackford, Demmel, Dongarra, Du Croz, Greenbaum, Hammarling, McKenney *et al*. 1999), all of above can be sped up automatically by Intel MKL (math kernel library) if it was linked with R by users, fast operations for dense and sparse matrix were implemented with the help of package **Matrix** (Douglas, Martin, Timothy, Jens, Jason, and R Core Team 2021), and only the main functions used for data and parameter input were written into pure R language, ensuring **hibayes** with a pretty higher computing efficiency.

For genotype loading, it is expensive to code it into numeric covariate matrix and read it into memory for each time of analysis, we provided an additional function to convert the genotype into numeric memory-mapping file by using the package **bigmemory** (Michael, John, Peter, and Charles 2019; Kane, Emerson, and Weston 2013), this only needs to be done at the first time, and no matter how big the number of individuals or markers in the genotype is, the memory-mapping file could be attached into memory on-the-fly within several minutes, making **hibayes** very promising in handing big data.

The package **hibayes** is available from CRAN at https://cran.r-project.org/package=hibayes. The latest version in development can be installed from GitHub at https://github.com/YinLiLin/hibayes. This article refers to version 1.0.1. The main available functions provided by **hibayes** are as following Table 2:

**Table 2:** Main functions provided by **hibayes**.

| Function name | Descriptions |
| --- | --- |
| `read_plink` | Converts genotype file in **PLINK** binary format into memory-mapping file format. |
| `ldmat` | Constructs variance-covariance matrix (**V**) of markers across the whole genome to fit summary level Bayesian model. |
| `bayes` | Fits individual level Bayesian model. |
| `sbayes` | Fits summary level Bayesian model. |
| `ssbayes` | Fits single-step Bayesian model. |

The detailed arguments for the functions in above table can be seen by typing the corresponding function name with a header of symbol ?” in R, for example, `>?bayes`.

## 4. Quick start with simple examples

In the following, we display some examples to run different Bayesian regression models using the tutorial data attached in **hibayes**, including the input file format, settings of main parameters, returned lists of results, as well as relevant visualizations of some important genetic parameters. We start by installing and loading **hibayes**:

~~~
*R> install.packages(“hibayes”)
R> library(“hibayes”)*
Loading required package: bigmemory
Loading required package: Matrix
Full description, Bug report, Suggestion and the latest codes:
https://github.com/YinLiLin/hibayes
~~~

### 4.1. Examples for individual level Bayesian model

To fit individual level Bayesian model, at least the phenotypic records (***n*** elements, missing should be marked as ‘NA’), numeric genotype (***n*** * ***m, n*** is the number of individuals, ***m*** is the number of genetic markers) should be provided. Users can load the phenotype and genotype data that coded by other software by the R function `read.table` to fit model, note that ‘NA’ is not allowed in genotype data,

~~~
*R> # load phenotype
R> pheno <- read.table(“your_pheno.txt”)
R> # load genotype
R> geno <- read.table(“your_geno.txt”)*
~~~

genotype should be coded in digits, either in c(0, 1, 2) or c(−1, 0, 1) is acceptable. Before fitting the model, it should be noted that the order of individuals should be exactly the same between phenotype and genotype, users should adjust it in prior, for example:

~~~
*R> geno.id <- read.table(“your_genoid.txt”)
R> # supposing the first column is the individual id
R> pheno <- pheno[match(geno.id[, 1], pheno[, 1]),]*
~~~

Additionally, we purposely provide a function `read_plink` in **hibayes** to load **PLINK** binary files into memory, and simultaneously construct memory-mapping file on local disk. For example, load the attached tutorial data in **hibayes**:

~~~
*R> bfile_path <- system.file(“extdata”, “example”, package = “hibayes”)
R> data <- read_plink(bfile = bfile_path, mode = “A”, threads = 4)*
~~~

the argument `bfile` is the prefix of binary files, `mode` can be set to “A” or “D” for additive and dominant genetic effect, respectively. In this function, missing genotype will be replaced by the major genotype of each allele. **hibayes** will code the genotype ***A***_1_***A***_1_ as **2, *A***_1_***A***_2_ as **1**, and ***A***_2_***A***_2_ as **0**, where ***A***_1_ is the first allele of each marker in *\*.bim* file, therefore, the estimated effect size is on ***A***_1_ allele, users should pay attention to it when a process involves marker effect. This data conversion only needs to be done at the first time, and no matter how big the number of individuals or markers in the genotype is, the memory-mapping file could be attached into memory on-the-fly within several minutes:

~~~
*R> geno <- attach.big.matrix(“./example.desc”)
R> map <- read.table(“./example.map”, header=TRUE)*
~~~

By default, the memory-mapped files are directed into the work directory, users could redirect to a new path by the argument `out` as follows:

~~~
*R> data <- read_plink(bfile = bfile_path, out=“./test”)*
~~~

then pick up the phenotype and genotype from the returned list, a quick view of the example data:

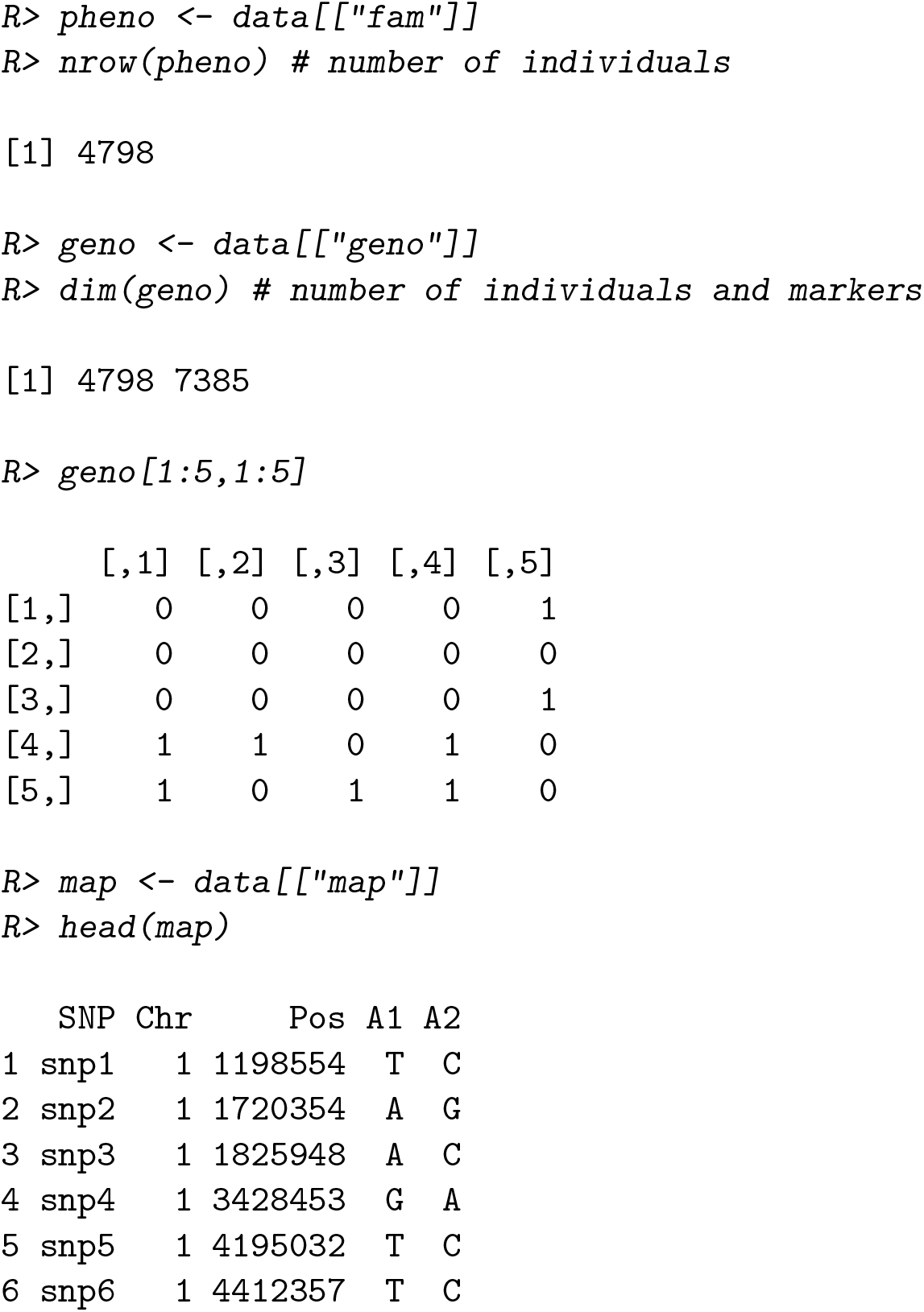

for the situation that the phenotype and genotype are loaded from binary files, we need not adjust the order of individuals additionally. Now, we can fit individual level Bayesian model as follows:

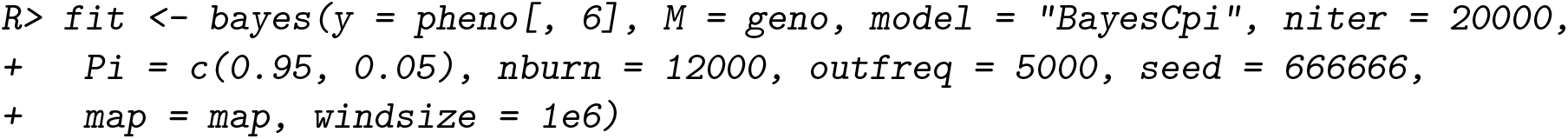
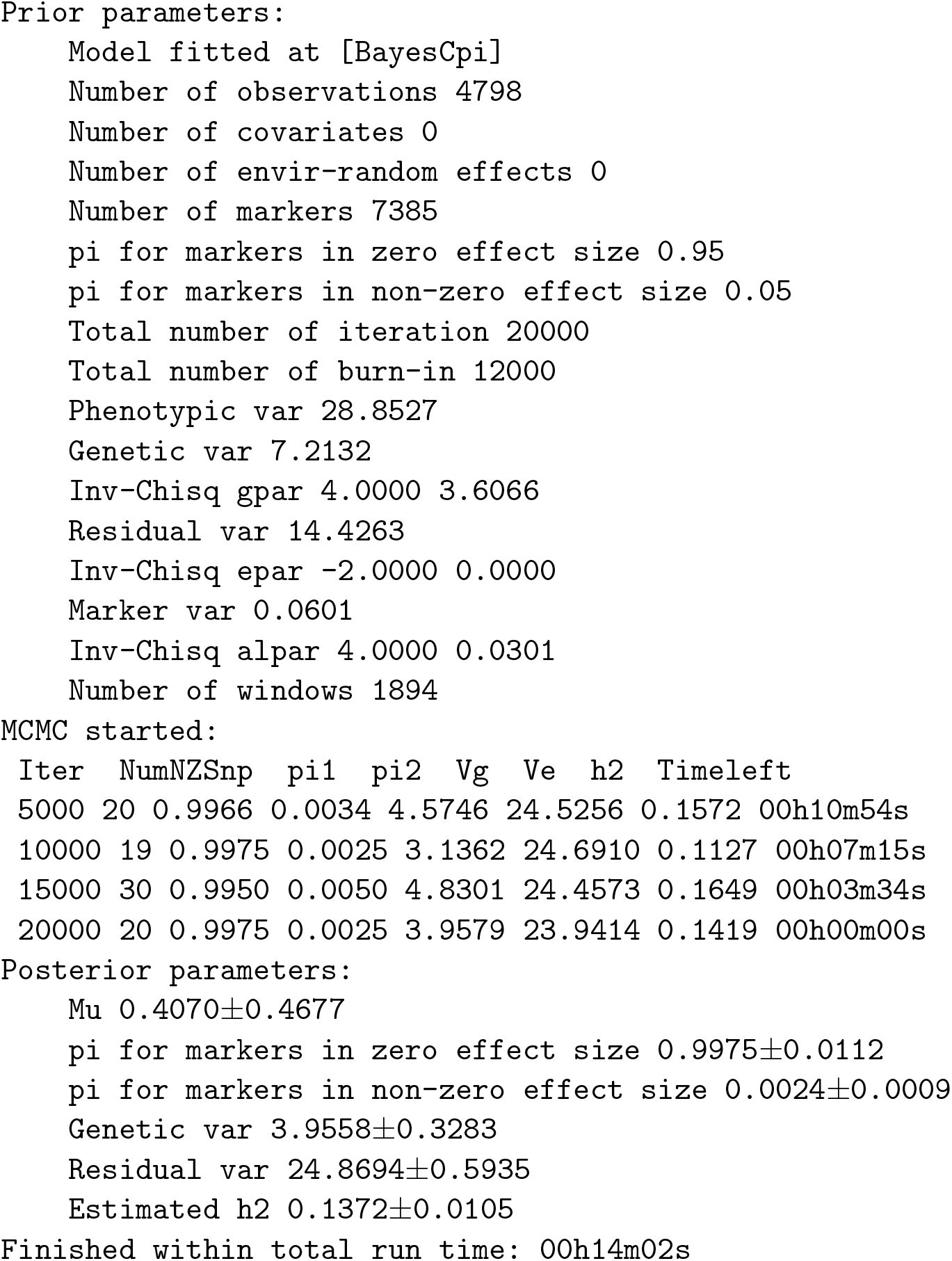

users can choose one of the methods in Table 1 by the argument `model`, change the total number of iterations and discarded number of iterations by the arguments `niter` and `nburn`, respectively. The printed log message recorded the descriptive information for input data and parameters, the convergence details of estimated parameters during the MCMC process, the time that remains for running, and summary statistics for some of main genetic parameters. Also, user can choose to turn off the LOG message by setting the argument `verbose = FALSE`.

It should be noted that the arguments `map` and `windsize` are optional, and they are only valid for GWAS analysis, and `windsize` is used to control the size of the windows, the number of markers in a window is not fixed. Alternatively, users can also choose another argument `windnum` (e.g., `windnum = 10`), which can be used to control the fixed number of markers in a window, the size for the window is not fixed for this case. Attentively, every marker should have clear physical position for the downstream genome cutting, however, if users are not going to run GWAS, these two arguments can be ignored.

The returned lists for the function `bayes` are

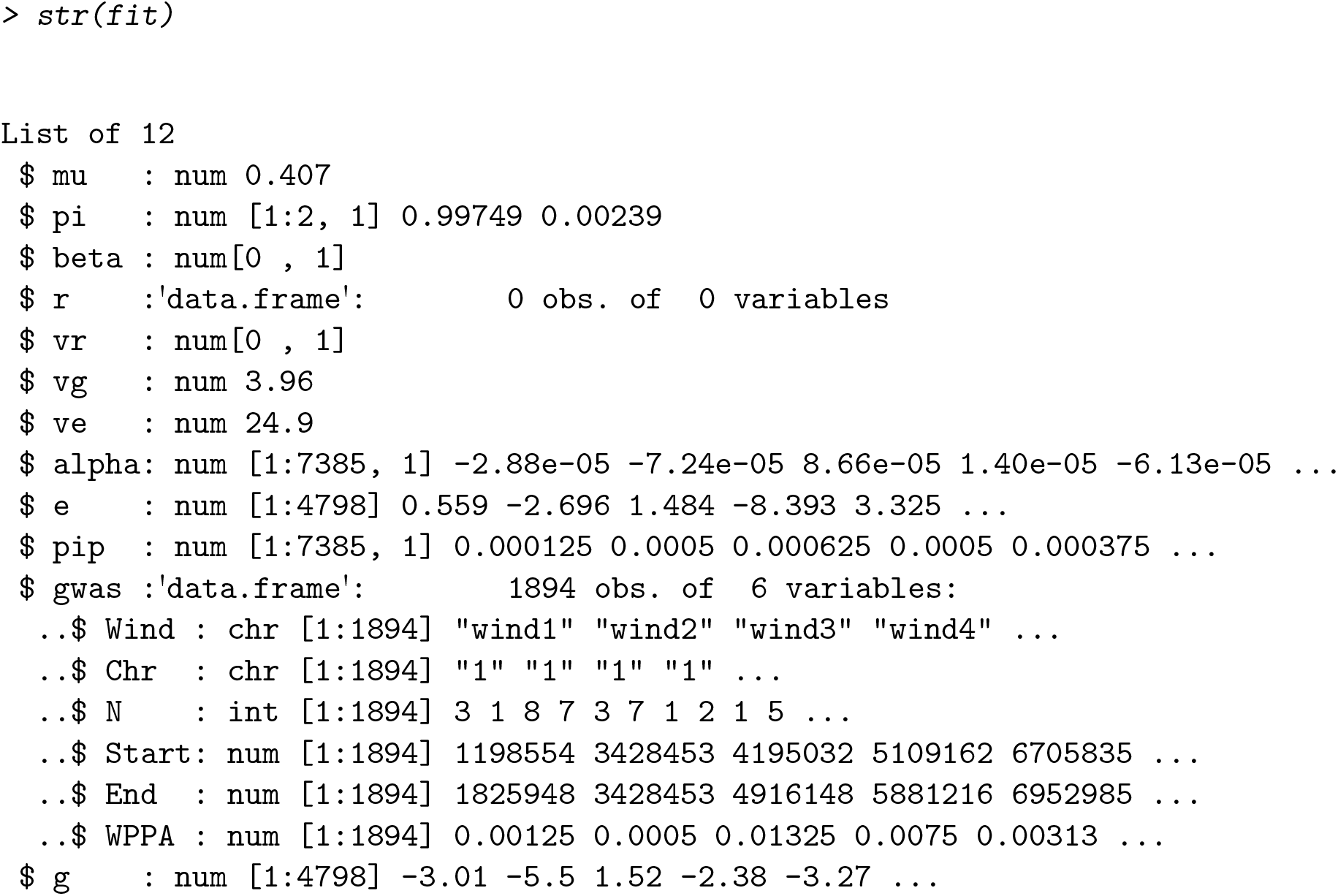

all the returned parameters estimated above are fully consistent with the Equation 2. As no fixed effects, covariates are specified for argument `X`, and also no environmental random effects for argument `R`, thus the returned lists `beta`, `r`, `vr`, are empty. For more details of how to model fixed effects, covariates, and environmental random effects, please refer to Section 4.3.1. The list `e` is the vector of residuals of the model, with the equal length and same order with the dependent variable ***y***. The list `gwas` is only visible when the the arguments `map` and `windsize` (or `windnum`) are specified by the users.

For genomic prediction, we can directly get the estimated effect size for all genetic markers, and the GEBVs for all individuals from the returns:

~~~
*R> SNPeffect <- fit[[“alpha”]]
R> gebv <- fit[[“g”]]*
~~~

we can visualize the marker effect size using the **CMplot** package (Yin 2021), as shown in the Manhattan plot (Figure 1):

**Figure 1:**
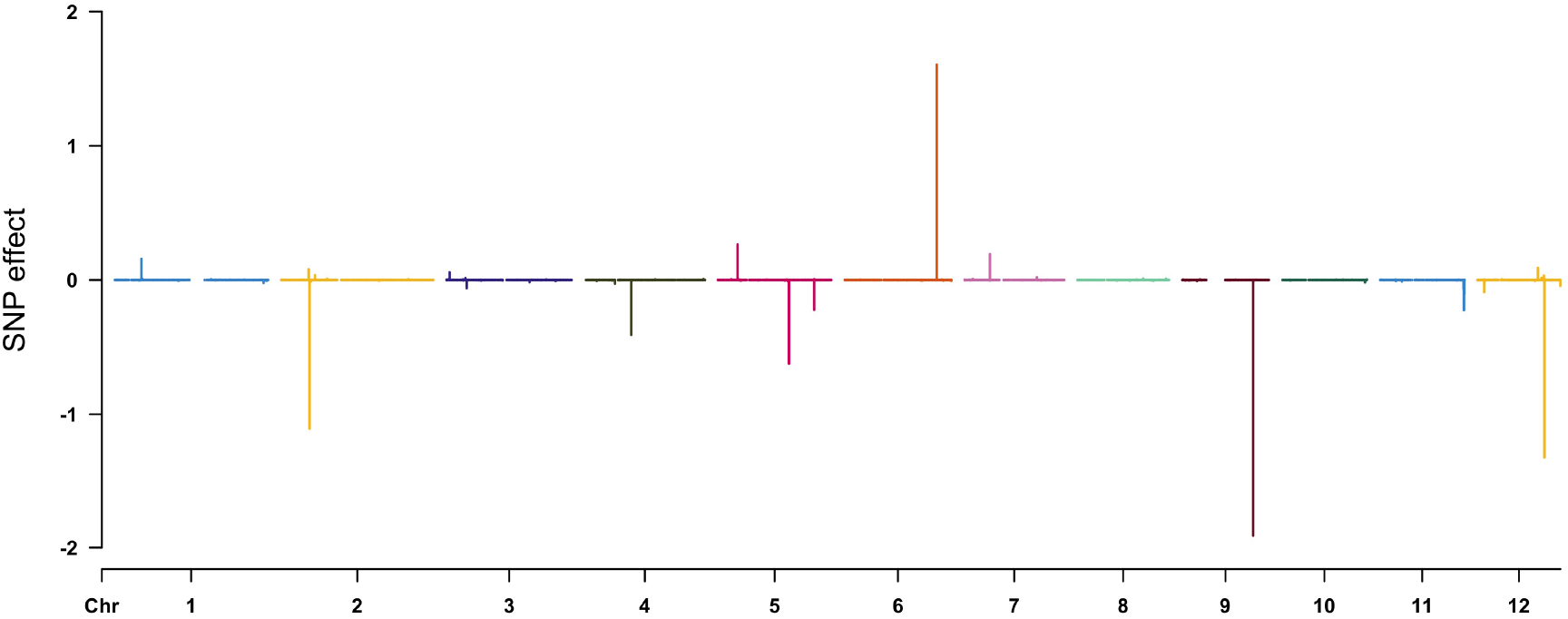
Estimated effect size of the genetic markers across the entire genome.

~~~
*R> library(CMplot)
R> CMplot(cbind(map[,1:3], SNPeffect), type = “h”, plot.type = “m”,
+ LOG10 = FALSE, ylab=“SNP effect”)*
~~~

we can also calculate and visualize the proportion of phenotypic variance explained (PVE) for each of genetic markers, as shown in the Figure 2:

**Figure 2:**
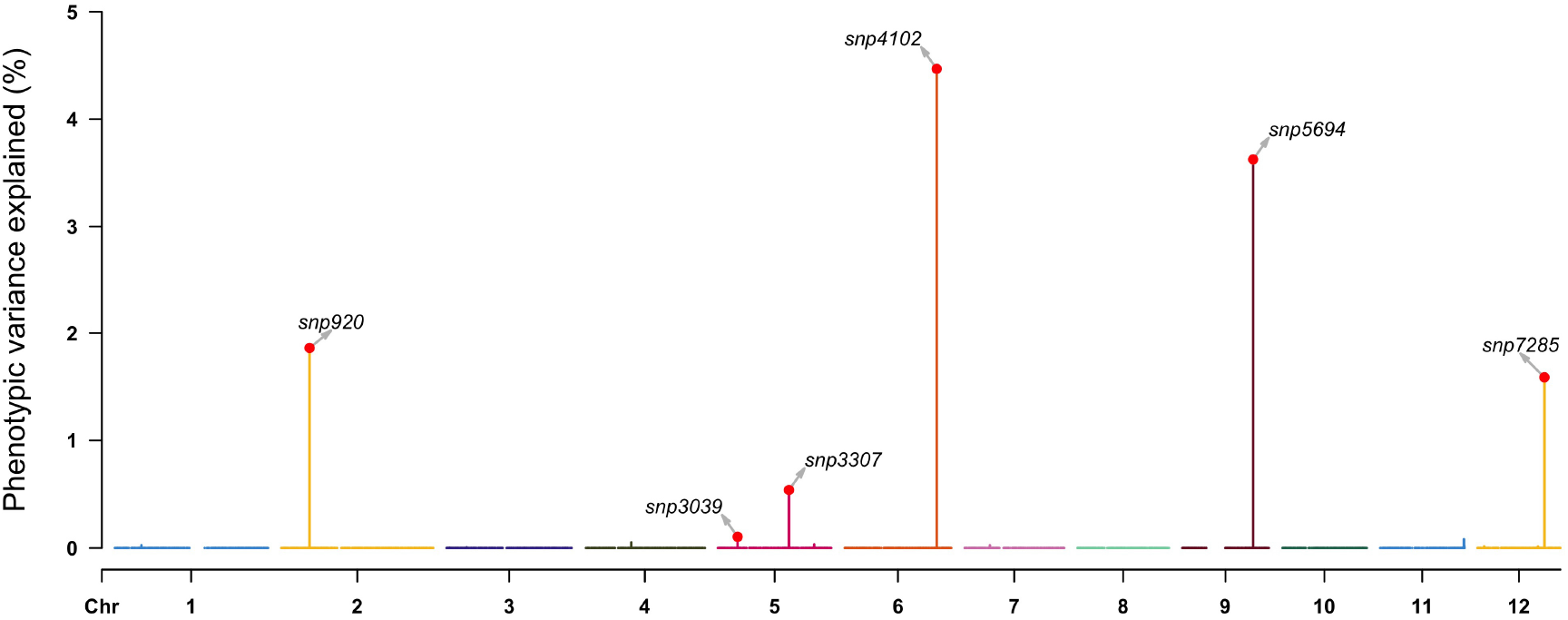
The proportion of explained phenotypic variance for each of markers.

~~~
*R> pve <- apply(as.matrix(geno), 2, var) * (fit[[“alpha”]]^2) / var(pheno[, 6])
R> highlight <- map[pve > 0.001,1]
R> CMplot(cbind(map[, 1:3], 100 * pve), type = “h”, plot.type = “m”,
+ LOG10 = FALSE, ylab = “Phenotypic variance explained (%)”,
+ highlight = highlight, highlight.text = highlight)*
~~~

For GWAS analysis, if the arguments `map` and `windsize` are detected in the input commands, **hibayes** will return a list named “gwas” as shown in following format:

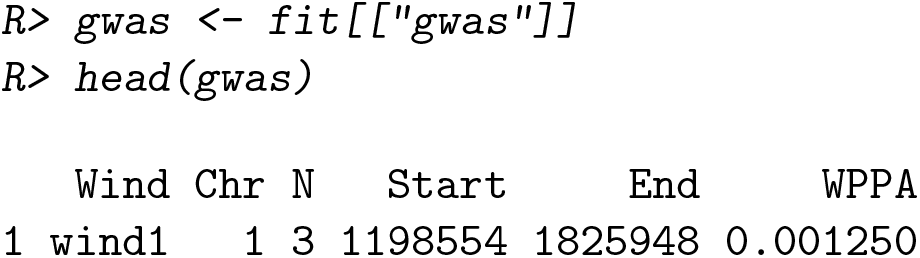
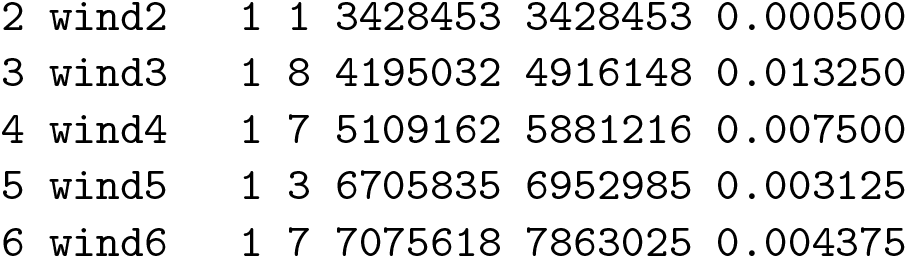

the first column records the names of all windows, second column is a vector of chromosomes, the third column reports the counted number of genetic markers in each of windows, the 4th and 5th columns are the physical position for the first and last genetic marker in each of windows respectively, and the last column lists the computed WPPA. To visualize WPPA as general Manhattan plot, we need to transform it into p-values as follows:

~~~
*R> highlight <- gwas[(1 - gwas[, “WPPA”]) < 0.01, 1]
R> CMplot(cbind(gwas[, c(1, 2, 4)], 1 - gwas[, “WPPA”]), type = “h”,
+ plot.type = “m”, LOG10 = TRUE, threshold = 0.01, ylim = c(0, 5),
+ ylab = expression(-log[10](1 - italic(WPPA))), highlight = highlight,
+ highlight.col = NULL, highlight.text = highlight)*
~~~

following Figure 3 visualizes the association results for all windows, the red line represents a significant level of 0.01, the windows that exceed this threshold are considered to be significantly associated with the trait of interest.

**Figure 3:**
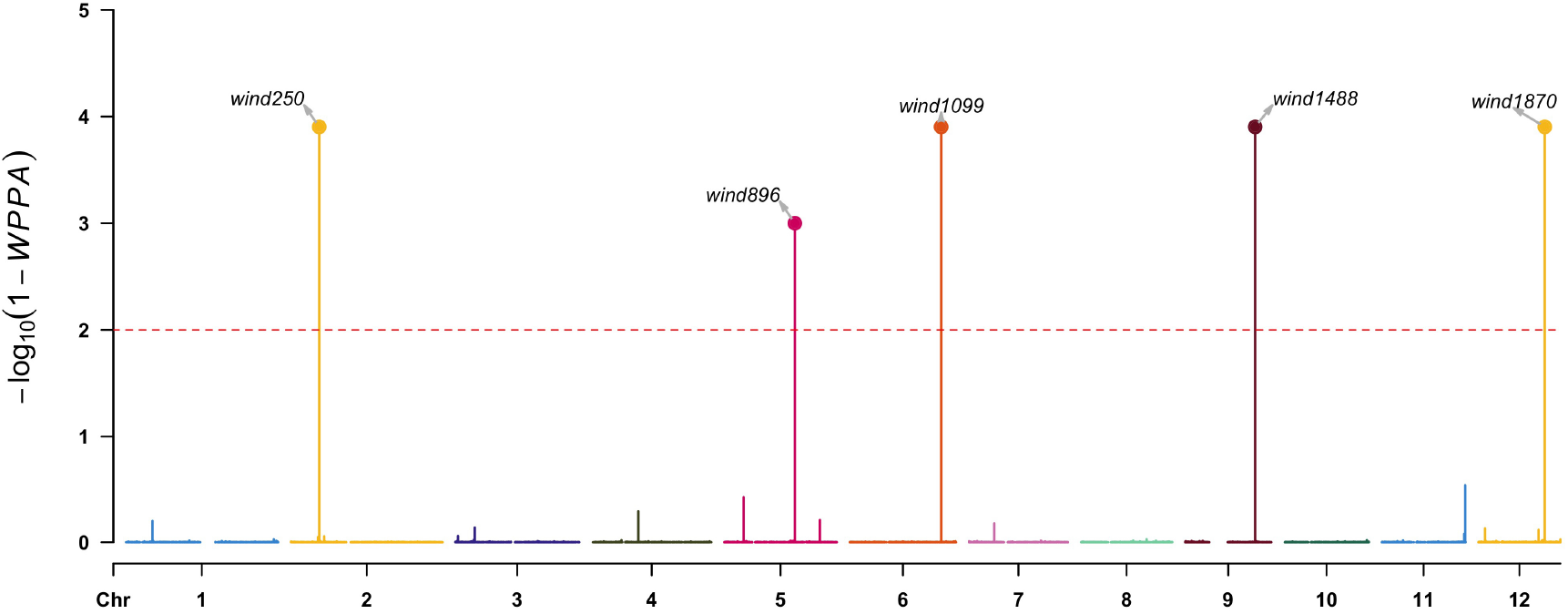
The derived window posterior probability of association from MCMC process.

However, it is still difficult to know which genetic markers in those significant windows are the causal variations, so we need to further explore the association significance for the markers. In **hibayes**, we also reported the posterior inclusive probability for every single marker, which could be used for an reference of importance to the trait, thus next step we can visualize PIP for a specific chromosome or region of interest as following Figure 4:

**Figure 4:**
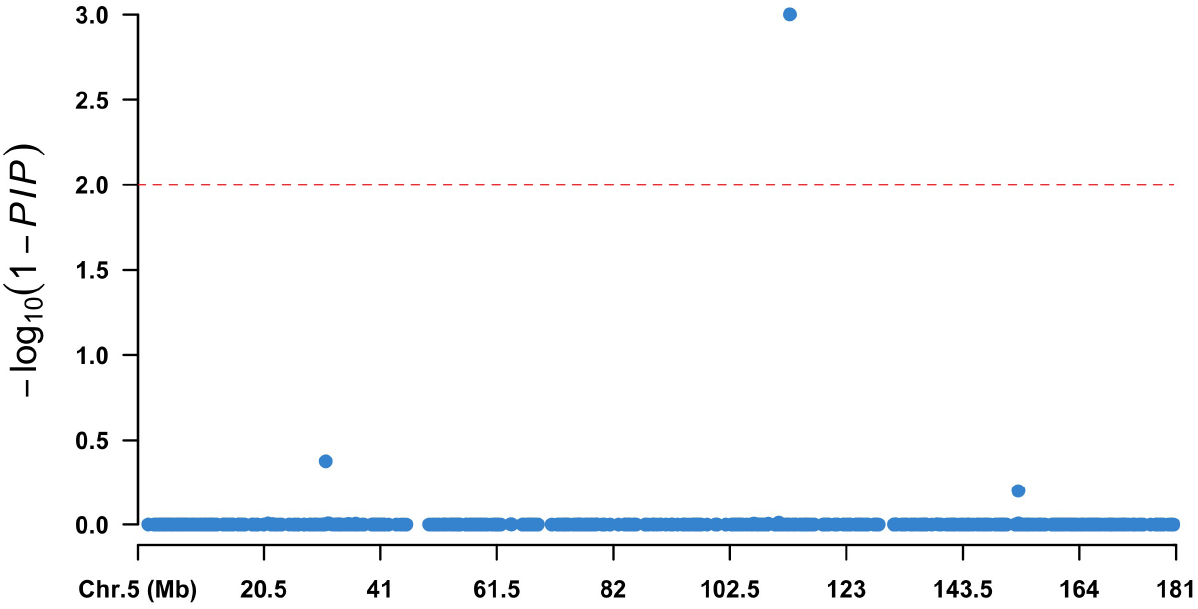
The posterior inclusive possibility for each of markers across Chromosome 5.

~~~
*R> data <- cbind(map[, 1:3], (1 - fit[[“pip”]]))
R> chr5 <- data[data[, 2] == 5,]
R> CMplot(chr5, plot.type = “m”, width = 9, height = 5, threshold = 0.01,
+ ylab = expression(-log[10](1 - italic(PIP))), LOG10 = TRUE,
+ amplify = FALSE)*
~~~

### 4.2. Examples for summary level Bayesian model

To fit summary level data based Bayesian model, the variance-covariance matrix calculated from the reference panel, and the summary data should be provided. The variance-covariance matrix (***V***) can be calculated by **hibayes** using either public reference genotype panel or personal genotype in hand. Taking the tutorial data attached in **hibayes** for an example:

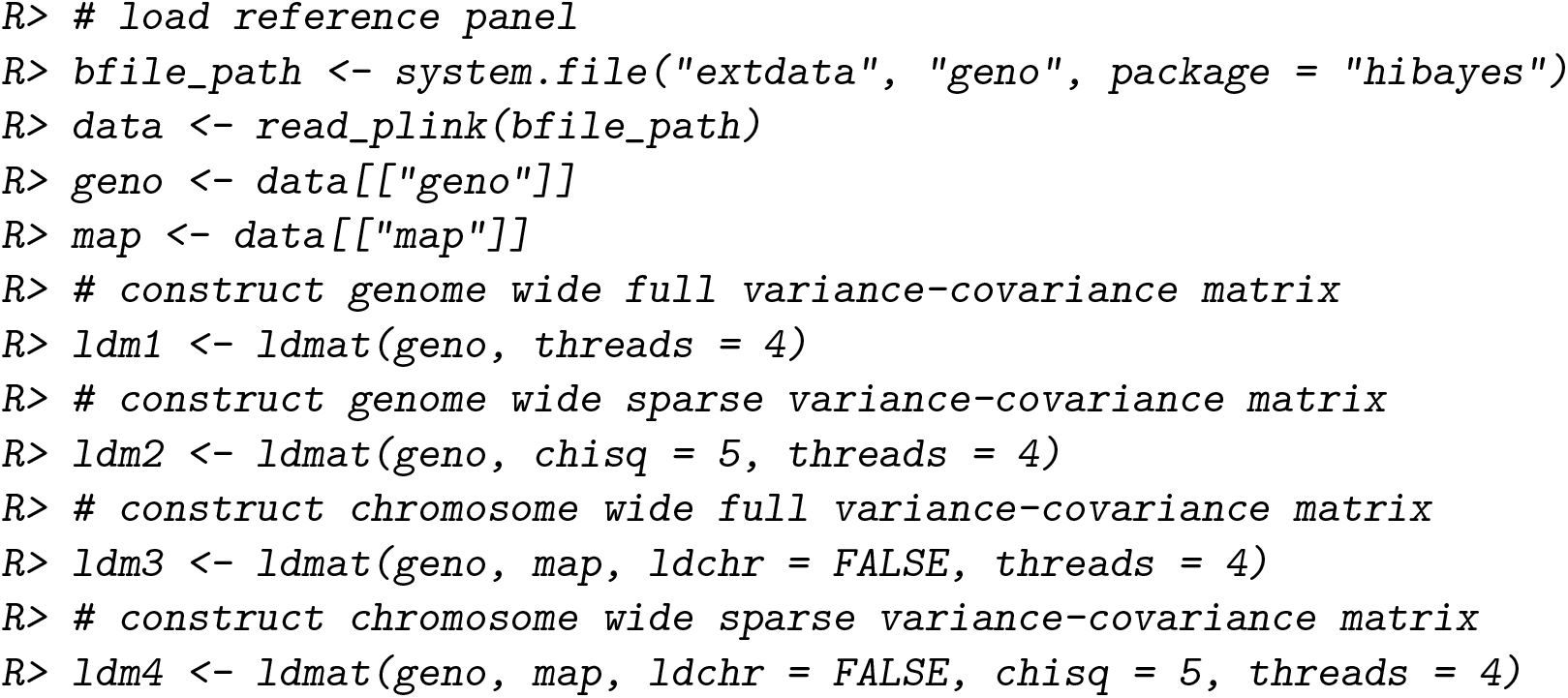

where the argument `chisq` is the chi-square threshold used for making sparse matrix, the degree of sparseness increases from “ldm1” to “ldm4”.

Load the tutorial summary data:

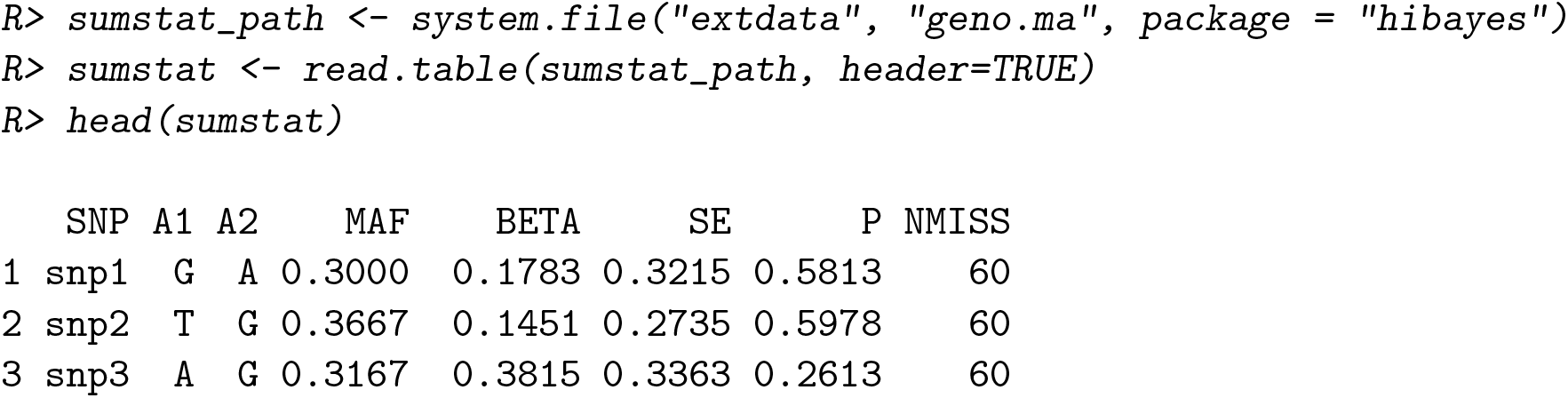
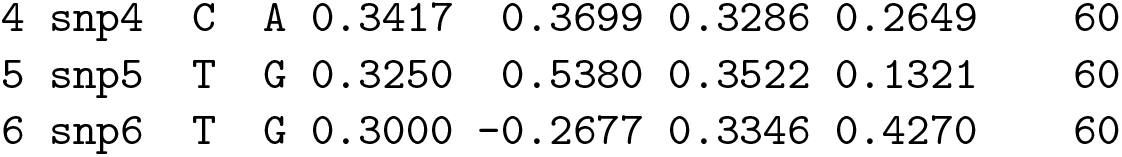

The order of genetic markers should be fully consistent between variance-covariance matrix and the summary data, thus prior adjustment on the order of genetic markers is required before fitting the model:

~~~
*R> sumstat <- sumstat[match(map[, 1], sumstat[, 1]),]*
~~~

Now we can fit summary level Bayesian model as follows:

~~~
*R> fit <- sbayes(sumstat = sumstat, ldm = ldm1, model = “BayesCpi”*,
*+ niter = 20000, Pi = c(0.95, 0.05), nburn = 12000, seed = 666666*,
*+ map = map, windsize=1e6)*
~~~

As described in Section 2.2, the returned lists are less than individual level Bayesian model:

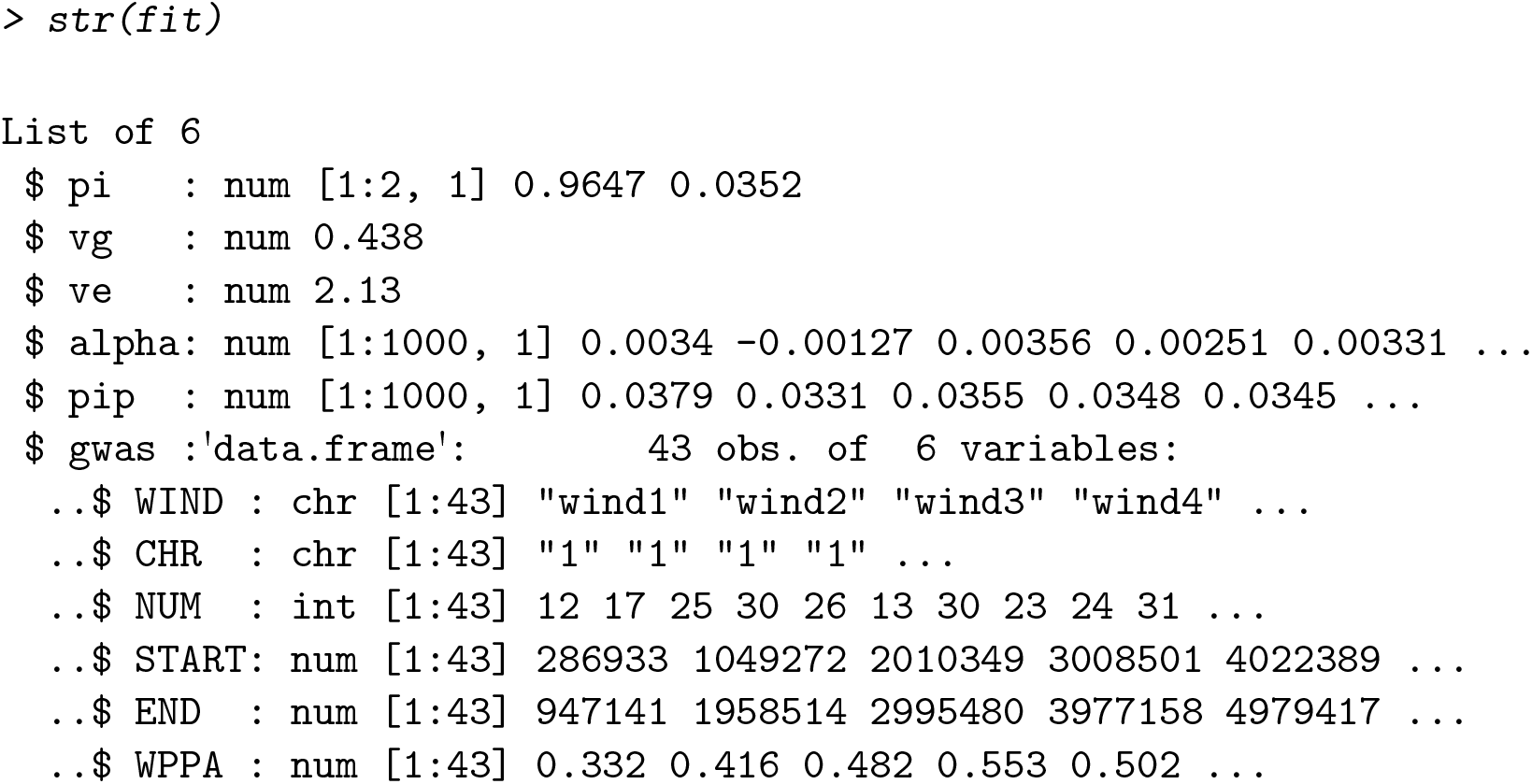

Similarly, we can visualize any of the estimated parameters for genomic prediction or genome-wide association studies as shown in Section 4.1

### 4.3. Examples for single-step Bayesian model

To fit single-step Bayesian model, at least the phenotype (***n***_1_, the number of individuals with phenotypic records), numeric genotype matrix (***n***_2_ * ***m***, where ***n***_2_ is the number of genotyped individuals, ***m*** is the number of markers), and pedigree information (***n***_3_ * **3**, the three columns are “id”, “sir”, “dam” orderly) should be provided, ***n***_1_, ***n***_2_, ***n***_3_ can be different, all the individuals in the pedigree will be predicted, including genotyped and non-genotyped, therefore the total number of predicted individuals depends on the number of unique individuals in pedigree. Taking the tutorial data attached in **hibayes** as an example:

loading and viewing the tutorial phenotype file,

|  | id | y | scale | group | sex |
| --- | --- | --- | --- | --- | --- |
| 1 | ind1 | -0.5796816 | 0.77 | g1 | m |
| 2 | ind2 | -2.0224628 | -1.02 | g2 | m |
| 3 | ind3 | -1.4807132 | 0.52 | g1 | f |
| 4 | ind4 | -3.0303065 | -1.05 | g4 | m |
| 5 | ind5 | 2.1881874 | 2.06 | g3 | m |
| 6 | ind6 | -3.2110719 | -1.94 | g4 | m |

loading and viewing the tutorial pedigree file,

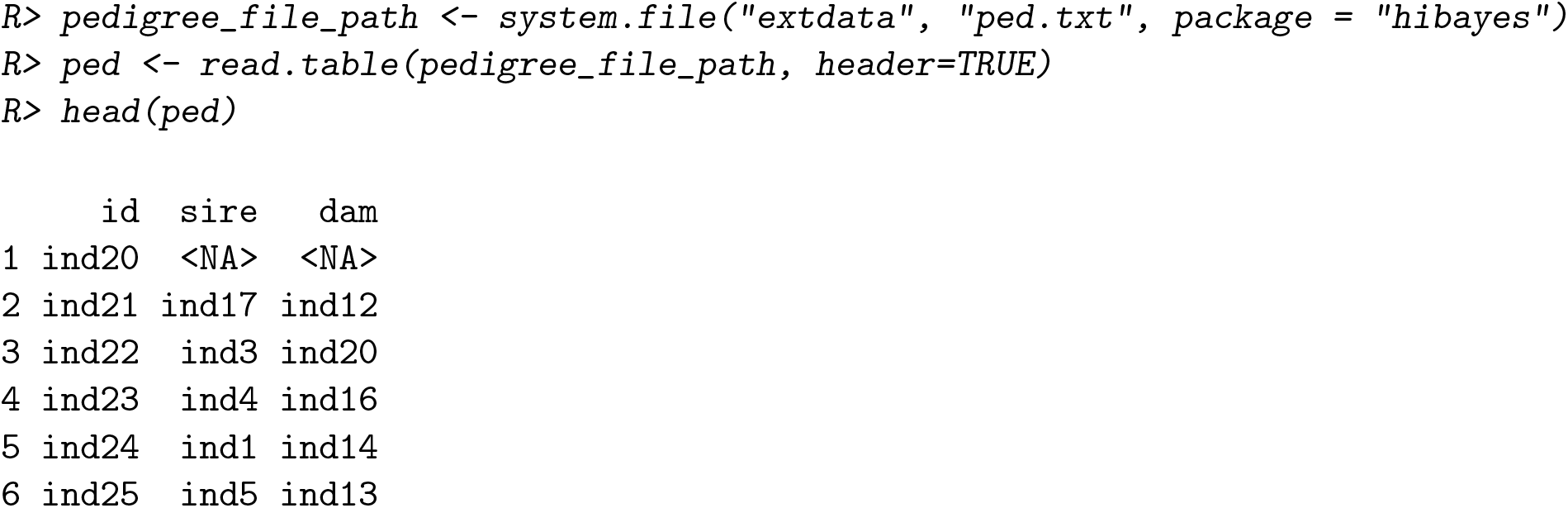

|  | id | sire | dam |
| --- | --- | --- | --- |
| 1 | ind20 | <NA> | <NA> |
| 2 | ind21 | ind17 | ind12 |
| 3 | ind22 | ind3 | ind20 |
| 4 | ind23 | ind4 | ind16 |
| 5 | ind24 | ind1 | ind14 |
| 6 | ind25 | ind5 | ind13 |

missing values in pedigree should be marked as “NA”, the columns must exactly follow “id”, “sir”, and “dam” in order.

converting and viewing the tutorial genotype data:

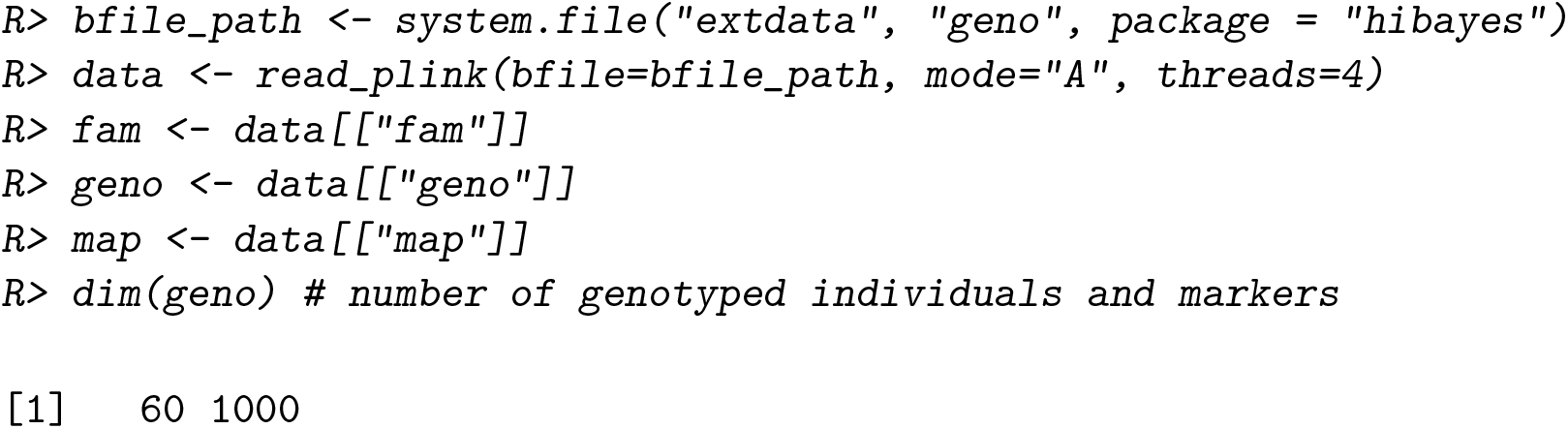

different with individual Bayesian model, it does not require the order of individuals in all data file to be consistent, **hibayes** can adjust the order automatically using the names provided by users:

~~~
*R> geno.id <- fam[, 2]
R> pheno.id <- pheno[, 1]*
~~~

#### Fixed effects, covariates and environmental random effects

For fixed effects, we can not fit it in the model directly for package **hibayes**, it should be converted in formula of designed model matrix, which can be achieved by the base function model.matrix.lm(…, na.action = “na.pass”) in R, for example, model “sex” as fixed effect, and “scale” as a covariate:

~~~
*R> X <- model.matrix.lm(∼ as.factor(sex) + as.numeric(scale)*,
*+ data = pheno, na.action = “na.pass”)
R> X <- X[, −1] #remove the intercept*
~~~

note that the attribute for fixed effects and covariates should be “factor” and “numeric” respectively.

For environmental random effects, there are no additional format transformations required, simply pick them out from the phenotype data, for example, model “group” as an environmental random effect:

~~~
*R> R <- pheno[, c(“group”)]*
~~~

Now we can fit single-step Bayesian model using all the data above:

~~~
*R> fit <- ssbayes(y = pheno[, 2], y.id = pheno.id, M = geno, M.id = geno.id*,
*+ P = ped, X = X, R = R, model = “BayesR”, niter = 20000, nburn = 12000*,
*+ Pi = c(0.95, 0.02, 0.02, 0.01), fold = c(0, 0.0001, 0.001, 0.01)*,
*+ outfreq = 500, seed = 666666, map = map, windsize = 1e6)*
~~~

the arguments y, `y.id`, `M`, `M.id`, `P` must be provided, other arguments are optional, if users want to implement GWAS analysis, the arguments `map`, `windsize` (or `windnum`) should be provided and specified. Compared with individual level Bayesian model, single-step Bayesian model returns two additional lists, `J` and `epsilon`, referring to Equation 26,

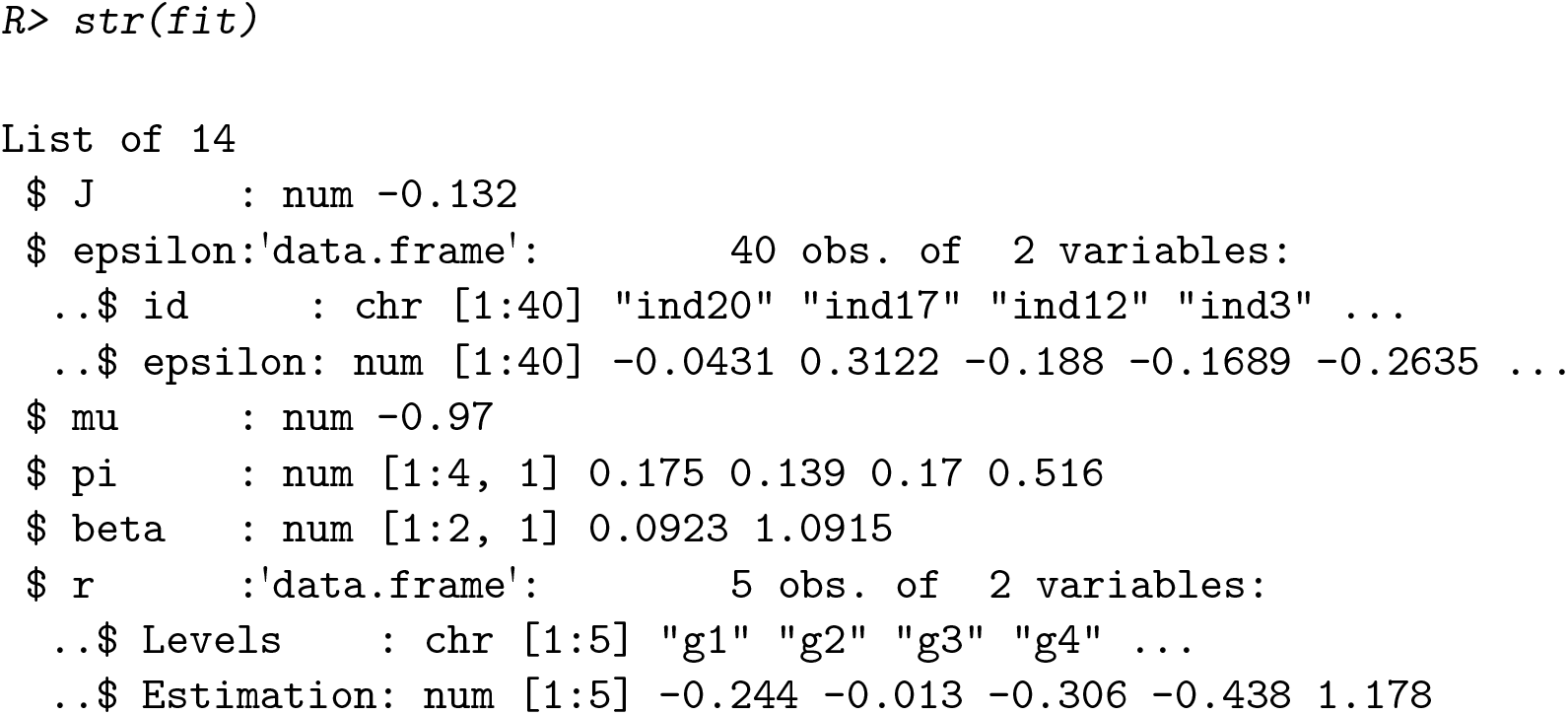

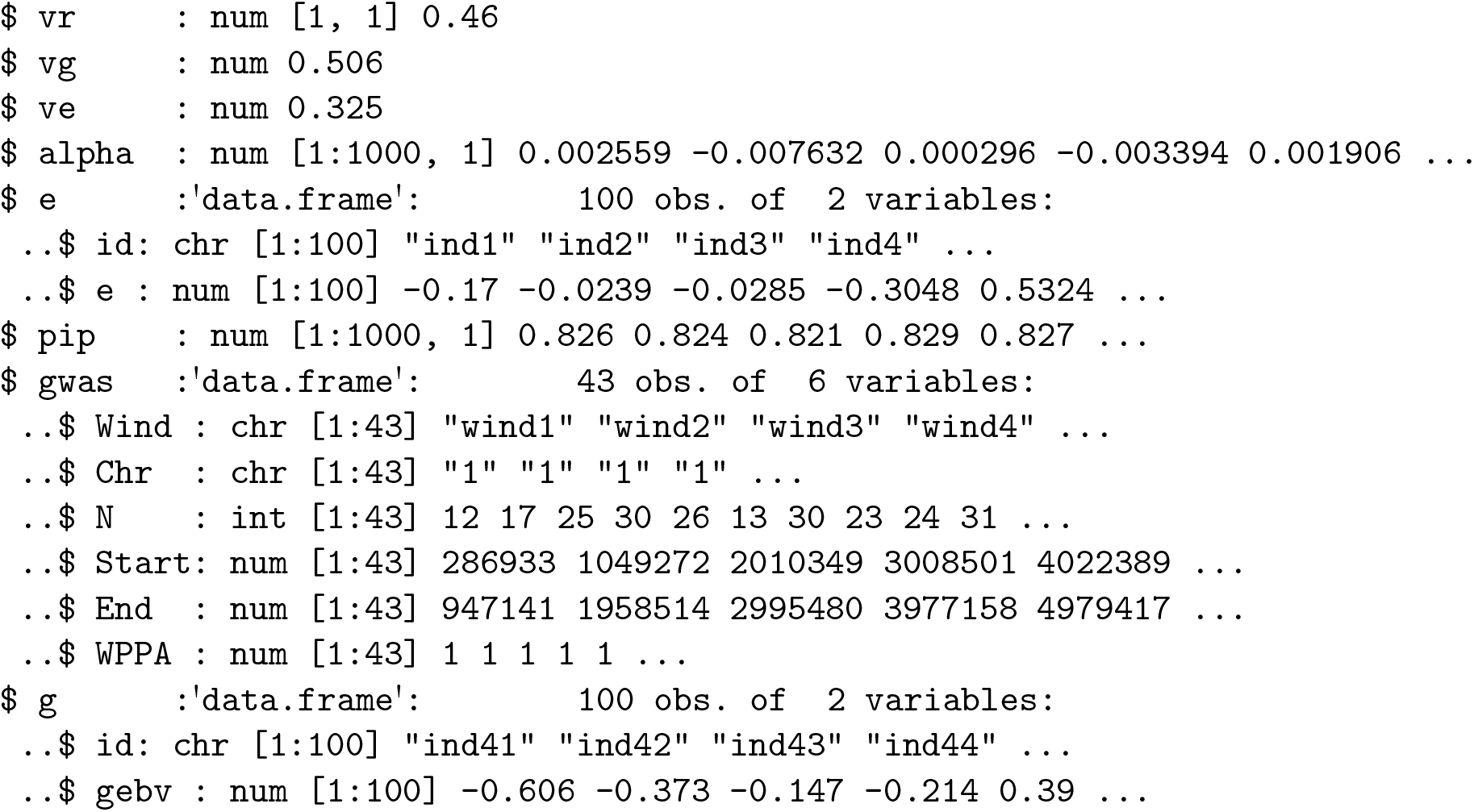

the returned list `g` is the vector of predicted GEBVs following the Equation 27 for all individuals in pedigree, the variance of imputation residuals and its standard deviation for non-genotyped individuals are printed at the end of LOG message, as it is generally not used for genetic evaluation, we do not attach it in the final returns.

## 5. Conclusion

The present paper is meant to provide a general overview on **hibayes**, the only one R package that can implement three types of Bayesian regression models with the richest methods achieved thus far. It is designed not only for genomic prediction, but also for genome-wide association studies. The package almost covers all the functions involved in genetic evaluation, including estimation of fixed effects and coefficients of covariates, environmental random effects and its corresponding variance, genetic and residual variance, heritability of traits, and effect size for all markers; computation of genomic estimated breeding values for both genotyped and non-genotyped individuals, phenotype/genetic variance explained for single or multiple markers; and derivation of posterior probability of association of the genomic window and posterior inclusive probability of markers. We roughly compared the inputs, direct returns, available methods and models, and relevant functionalities for the most widely used tools in fitting Bayesian regression models, as shown in Table 3, **hibayes** is more comprehensive over other tools for genomic analysis.

**Table 3:** Comparisons of inputs, direct, returns, and available models and methods for **BGLR, JWAS, GCTB**, and **hibayes**.

| Models | Functionalities | BGLR | JWAS | GCTB | hibayes |
| --- | --- | --- | --- | --- | --- |
|  | Language | R, C++ | Julia | C++ | R, C++ |
|  | Version | 1.0.9 | 1.0.0 | 2.0.2 | 1.0.1 |
| **Individual level Bayesian model** | Covariates | ✓ | ✓ | ✓ | ✓ |
|  | Fixed effects | ✓ | ✓ | × | ✓ |
|  | Random effects | × | ✓ | × | ✓ |
|  | Environmental interaction | ✓ | ✓ | × | ✓ |
|  | Variance components | ✓ | ✓ | ✓ | ✓ |
|  | PIP | × | ✓ | ✓ | ✓ |
|  | WPPA | × | ✓ | × | ✓ |
|  | Marker effects | ✓ | ✓ | ✓ | ✓ |
|  | GEBVs | × | ✓ | × | ✓ |
|  | Residuals | × | × | × | ✓ |
|  | Achieved methods | RR, A, B, Cpi, L | RR, A, B, Bpi, C, Cpi, L | RR, A, B, Bpi, C, Cpi, S, R | RR, A, B, Bpi, C, Cpi, L, BSLMM, R |
| **Summary level Bayesian model** | Variance components | × | × | ✓ | ✓ |
|  | PIP | × | × | ✓ | ✓ |
|  | WPPA | × | × | × | ✓ |
|  | Marker effects | × | × | ✓ | ✓ |
|  | Achieved methods | × | × | C, Cpi, R | CG, RR, A, B, Bpi, C, Cpi, L, R |
| **Single-step Bayesian model** | Covariates | × | ✓ | × | ✓ |
|  | Fixed effects | × | ✓ | × | ✓ |
|  | Random effects | × | ✓ | × | ✓ |
|  | Environmental interaction | × | ✓ | × | ✓ |
|  | Variance components | × | ✓ | × | ✓ |
|  | PIP | × | ✓ | × | ✓ |
|  | WPPA | × | ✓ | × | ✓ |
|  | Marker effects | × | ✓ | × | ✓ |
|  | GEBVs | × | ✓ | × | ✓ |
|  | Residuals | × | × | × | ✓ |
|  | Achieved methods | × | RR, A, B, Bpi, C, Cpi, L | × | RR, A, B, Bpi, C, Cpi, L, R |

The arguments of main functions and the alias of returns in **hibayes** are highly consistent with the equations listed in this paper, and the functional style and idiomatic implementation in R make the package easy to use, flexible to extend, and transparent to validate. Although only a small selection of the modeling options available in **hibayes** are discussed in detail, we hope that this article can serve as a good starting point to further explore the capabilities of the package.

For the future, we have several plans on how to improve the functionality of **hibayes**. We will keep on updating **hibayes** with the latest advanced Bayesian models and methods of broadly interest in the domain of genomic prediction or genome-wide association studies, making **hibayes** always be fresh to the users or academic researchers. Also, we will next achieve multiple traits Bayesian regression models in **hibayes**, which can be used to estimate genetic correlation among traits. Furthermore, the MCMC process requires tremendous amounts of time to reach the convergence, how to partly speed up MCMC by parallel computing would be our next focus of work on **hibayes**.

## Acknowledgments

First of all, we would like to thank Dr. Jian Zeng from the University of Queensland for his kindly guidance in developing summary data based Bayesian regression model in **hibayes**. We want to thank Prof. Hao Cheng from the University of California, Davis, for his nice package **JWAS** written in Julia, which helps a lot in making comparisons for developing parts of methods of single-step Bayesian model in **hibayes**. We sincerely thank Thomas A. Gavin, Professor Emeritus, Cornell University, for help with editing this paper. Furthermore, We also would like to sincerely thank the many users who reported bugs or valuable ideas for new features, thus helping to continuously improvement for **hibayes**.

This work was supported by the National Natural Science Foundation of China [32102516, 32072725]; Science Foundation for Creative Research Groups of the Natural Science Foundation of Hubei Province [2020CFA006]; Cooperation funding of Huazhong Agricultural University - Shenzhen Institute of Agricultural Genomics, Chinese Academy of Agricultural Sciences; and the National Swine Industry Technology System [CARS-35].

## Notes

### Competing Interest Statement

The authors have declared no competing interest.

### Summary of Updates

We correct the functionalities of the compared software for Table 3 in the manuscript.

## References

Aguilar I, Misztal I, Johnson D, Legarra A, Tsuruta S, Lawlor T (2010). “Hot topic: A unified approach to utilize phenotypic, full pedigree, and genomic information for genetic evaluation of Holstein final score.” Journal of Dairy Science, 93(2), 743–752. doi:10.3168/jds.2009-2730.

Aliloo H, Pryce J, González-Recio O, Cocks B, Goddard M, Hayes B (2017). “Including non-additive genetic effects in mating programs to maximize dairy farm profitability.” Journal of Dairy Science, 100(2), 1203–1222. doi:10.3168/jds.2016-11261.

Anderson E, Bai Z, Bischof C, Blackford LS, Demmel J, Dongarra J, Du Croz J, Greenbaum A, Hammarling S, McKenney A, et al. (1999). LAPACK Users’ guide. SIAM. doi:10.1137/1.9780898719604.

Bulik-Sullivan BK, Loh PR, Finucane HK, Ripke S, Yang J, Patterson N, Daly MJ, Price AL, Neale BM (2015). “LD Score regression distinguishes confounding from polygenicity in genome-wide association studies.” Nature Genetics, 47(3), 291–295. doi:10.1038/ng.3211. URL https://github.com/bulik/ldsc.

Cheng H, Fernando R, Garrick D (2018). “JWAS: Julia implementation of whole-genome analysis software.” In Proceedings of the world congress on genetics applied to livestock production, volume 11, p. 859. URL https://reworkhow.github.io/JWAS.jl/latest/.

Christensen OF, Lund MS (2010). “Genomic prediction when some animals are not genotyped.” Genetics Selection Evolution, 42(1). doi:10.1186/1297-9686-42-2.

Dagum L, Menon R (1998). “OpenMP: an industry standard API for shared-memory programming.” IEEE Computational Science and Engineering, 5(1), 46–55. doi:10.1109/99.660313.

de los Campos G, Hickey JM, Pong-Wong R, Daetwyler HD, Calus MPL (2013). “Whole-Genome Regression and Prediction Methods Applied to Plant and Animal Breeding.” Genetics, 193(2), 327–345. doi:10.1534/genetics.112.143313.

Douglas B, Martin M, Timothy AD, Jens O, Jason R, R Core Team (2021). “Matrix: Sparse and Dense Matrix Classes and Methods.” R package version 1.4-0, URL https://CRAN.R-project.org/package=Matrix.

Eddelbuettel D, François R (2011). “Rcpp: Seamless R and C++ integration.” Journal of Statistical Software, 40(8). doi:10.18637/jss.v040.i08. URL https://CRAN.R-project.org/package=Rcpp.

Eddelbuettel D, Sanderson C (2014). “RcppArmadillo: Accelerating R with highperformance C++ linear algebra.” Computational Statistics & Data Analysis, 71, 1054–1063. doi:10.1016/j.csda.2013.02.005. URL https://CRAN.R-project.org/package=RcppArmadillo.

Erbe M, Hayes B, Matukumalli L, Goswami S, Bowman P, Reich C, Mason B, Goddard M (2012). “Improving accuracy of genomic predictions within and between dairy cattle breeds with imputed high-density single nucleotide polymorphism panels.” Journal of Dairy Science, 95(7), 4114–4129. doi:10.3168/jds.2011-5019.

Fan B, Onteru SK, D. ZQ, Garrick DJ, Stalder KJ, Rothschild MF (2011). “Genome-Wide Association Study Identifies Loci for Body Composition and Structural Soundness Traits in Pigs.” PLoS ONE, 6(2), e14726. doi:10.1371/journal.pone.0014726.

Fernando R, Toosi A, Wolc A, Garrick D, Dekkers J (2017). “Application of Whole-Genome Prediction Methods for Genome-Wide Association Studies: A Bayesian Approach.” Journal of Agricultural, Biological and Environmental Statistics, 22(2), 172–193. doi:10.1007/s13253-017-0277-6.

Fernando RL, Cheng H, Golden BL, Garrick DJ (2016). “Computational strategies for alternative single-step Bayesian regression models with large numbers of genotyped and non-genotyped animals.” Genetics Selection Evolution, 48(1). doi:10.1186/s12711-016-0273-2.

Fernando RL, Dekkers JC, Garrick DJ (2014). “A class of Bayesian methods to combine large numbers of genotyped and non-genotyped animals for whole-genome analyses.” Genetics Selection Evolution, 46(1), 50. doi:10.1186/1297-9686-46-50.

García-Cortés L, Sorensen D (1996). “On a multivariate implementation of the Gibbs sampler.” Genetics Selection Evolution, 28(1), 121. doi:10.1186/1297-9686-28-1-121.

Habier D, Fernando RL, Kizilkaya K, Garrick DJ (2011). “Extension of the bayesian alphabet for genomic selection.” BMC Bioinformatics, 12(1). doi:10.1186/1471-2105-12-186.

Henderson CR (1975). “Best Linear Unbiased Estimation and Prediction under a Selection Model.” Biometrics, 31(2), 423. doi:10.2307/2529430.

Henderson CR (1976). “A Simple Method for Computing the Inverse of a Numerator Relationship Matrix Used in Prediction of Breeding Values.” Biometrics, 32(1), 69. doi:10.2307/2529339.

Kane MJ, Emerson J, Weston S (2013). “Scalable Strategies for Computing with Massive Data.” Journal of Statistical Software, 55(14). doi:10.18637/jss.v055.i14. URL https://doi.org/10.18637%2Fjss.v055.i14.

Lloyd-Jones LR, Zeng J, Sidorenko J, Yengo L, Moser G, Kemper KE, Wang H, Zheng Z, Magi R, Esko T, Metspalu A, Wray NR, Goddard ME, Yang J, Visscher PM (2019). “Improved polygenic prediction by Bayesian multiple regression on summary statistics.” doi:10.1101/522961.

Lund MS, Jensen CS (1999). “Blocking Gibbs sampling in the mixed inheritance model using graph theory.” Genetics Selection Evolution, 31(1), 3–24. doi:10.1186/1297-9686-31-1-3.

Madsen P, Sørensen P, Su G, Damgaard L, Thomsen H, Labouriau R, et al. (2006). “DMU-a package for analyzing multivariate mixed models.” In 8th World Congress on Genetics Applied to Livestock Production, volume 247. Belo Horizonte. URL https://dmu.ghpc.au.dk/dmu/DMU/.

Meuwissen THE, Hayes BJ, Goddard ME (2001). “Prediction of Total Genetic Value Using Genome-Wide Dense Marker Maps.” Genetics, 157(4), 1819–1829. doi:10.1093/genetics/157.4.1819.

Michael JK, John WE, Peter H, Charles DJ (2019). “bigmemory: Manage Massive Matrices with Shared Memory and Memory-Mapped Files.” R package version 4.5.36, URL https://CRAN.R-project.org/package=bigmemory.

Misztal I, Tsuruta S, Strabel T, Auvray B, Druet T, Lee D, et al. (2002). “BLUPF90 and related programs (BGF90).” In Proceedings of the 7th world congress on genetics applied to livestock production, volume 28. Montpellier. URL http://nce.ads.uga.edu/html/projects/programs/.

Moser G, Lee SH, Hayes BJ, Goddard ME, Wray NR, Visscher PM (2015). “Simultaneous Discovery, Estimation and Prediction Analysis of Complex Traits Using a Bayesian Mixture Model.” PLOS Genetics, 11(4), e1004969. doi:10.1371/journal.pgen.1004969.

Ozaki K, Ohnishi Y, Iida A, Sekine A, Yamada R, Tsunoda T, Sato H, Sato H, Hori M, Nakamura Y, Tanaka T (2002). “Functional SNPs in the lymphotoxin-gene that are associated with susceptibility to myocardial infarction.” Nature Genetics, 32(4), 650–654. doi:10.1038/ng1047.

Pérez P, de los Campos G (2014). “Genome-Wide Regression and Prediction with the BGLR Statistical Package.” Genetics, 198(2), 483–495. doi:10.1534/genetics.114.164442. URL https://CRAN.R-project.org/package=BGLR.

Purcell S, Neale B, Todd-Brown K, Thomas L, Ferreira MA, Bender D, Maller J, Sklar P, de Bakker PI, Daly MJ, Sham PC (2007). “PLINK: A Tool Set for Whole-Genome Association and Population-Based Linkage Analyses.” The American Journal of Human Genetics, 81(3), 559–575. doi:10.1086/519795. URL https://www.cog-genomics.org/plink/.

R Core Team (2013). “R: A language and environment for statistical computing.” URL https://www.R-project.org/.

Robinson MR, Kleinman A, Graff M, Vinkhuyzen AA, Couper D, Miller MB, Peyrot WJ, Abdellaoui A, Zietsch BP, Nolte IM, et al. (2017). “Genetic evidence of assortative mating in humans.” Nature Human Behaviour, 1(1), 1–13. doi:10.1038/s41562-016-0016.

Sorensen D, Gianola D (2002). Likelihood, Bayesian, and MCMC Methods in Quantitative Genetics. Springer New York. doi:10.1007/b98952.

VanRaden P (2008). “Efficient Methods to Compute Genomic Predictions.” Journal of Dairy Science, 91(11), 4414–4423. doi:10.3168/jds.2007-0980.

Vilhjálmsson BJ, Yang J, Finucane HK, Gusev A, Lindström S, Ripke S, Genovese G, Loh PR, Bhatia G, Do R, et al. (2015). “Modeling linkage disequilibrium increases accuracy of polygenic risk scores.” The american journal of human genetics, 97(4), 576–592. doi:10.1016/j.ajhg.2015.09.001.

Wray NR, Wijmenga C, Sullivan PF, Yang J, Visscher PM (2018). “Common Disease Is More Complex Than Implied by the Core Gene Omnigenic Model.” Cell, 173(7), 1573–1580. doi:10.1016/j.cell.2018.05.051.

Yang J,, Ferreira T, Morris AP, Medland SE, Madden PAF, Heath AC, Martin NG, Montgomery GW, Weedon MN, Loos RJ, Frayling TM, McCarthy MI, Hirschhorn JN, Goddard ME, and PMV (2012). “Conditional and joint multiple-SNP analysis of GWAS summary statistics identifies additional variants influencing complex traits.” Nature Genetics, 44(4), 369–375. doi:10.1038/ng.2213.

Yang J, Lee SH, Goddard ME, Visscher PM (2011). “GCTA: A Tool for Genome-wide Complex Trait Analysis.” The American Journal of Human Genetics, 88(1), 76–82. doi:10.1016/j.ajhg.2010.11.011. URL https://yanglab.westlake.edu.cn/software/gcta/#Overview.

Yi N, George V, Allison DB (2003). “Stochastic Search Variable Selection for Identifying Multiple Quantitative Trait Loci.” Genetics, 164(3), 1129–1138. doi:10.1093/genetics/164.3.1129.

Yi N, Xu S (2008). “Bayesian LASSO for Quantitative Trait Loci Mapping.” Genetics, 179(2), 1045–1055. doi:10.1534/genetics.107.085589.

Yin L (2021). “CMplot: Circle Manhattan Plot.” R package version 3.7.0, URL https://CRAN.R-project.org/package=CMplot.

Yin L, Zhang H, Li X, Zhao S, Liu X (2021a). “hibayes: Individual-Level, Summary-Level and Single-Step Bayesian Regression Model for Genome-Wide Association and Genomic Prediction.” R package version 1.0.0, URL https://CRAN.R-project.org/package=hibayes.

Yin L, Zhang H, Tang Z, Xu J, Yin D, Zhang Z, Yuan X, Zhu M, Zhao S, Li X, Liu X (2021b). “rMVP: A Memory-efficient, Visualization-enhanced, and Parallel-accelerated tool for Genome-Wide Association Study.” Genomics, Proteomics & Bioinformatics. doi:10.1016/j.gpb.2020.10.007. URL https://CRAN.R-project.org/package=rMVP.

Yin L, Zhang H, Zhou X, Yuan X, Zhao S, Li X, Liu X (2020). “KAML: improving genomic prediction accuracy of complex traits using machine learning determined parameters.” Genome Biology, 21(1). doi:10.1186/s13059-020-02052-w.

Zeng J, de Vlaming R, Wu Y, Robinson MR, Lloyd-Jones LR, Yengo L, Yap CX, Xue A, Sidorenko J, McRae AF, Powell JE, Montgomery GW, Metspalu A, Esko T, Gibson G, Wray NR, Visscher PM, Yang J (2018). “Signatures of negative selection in the genetic architecture of human complex traits.” Nature Genetics, 50(5), 746–753. doi:10.1038/s41588-018-0101-4. URL https://cnsgenomics.com/software/gctb/#Download.

Zhou X, Carbonetto P, Stephens M (2013). “Polygenic Modeling with Bayesian Sparse Linear Mixed Models.” PLoS Genetics, 9(2), e1003264. doi:10.1371/journal.pgen.1003264.

Zhou X, Stephens M (2012). “Genome-wide efficient mixed-model analysis for association studies.” Nature Genetics, 44(7), 821–824. doi:10.1038/ng.2310. URL https://github.com/genetics-statistics/GEMMA.

Zhu X, Stephens M (2016). “Bayesian large-scale multiple regression with summary statistics from genome-wide association studies.” doi:10.1101/042457. URL https://doi.org/10.1101%2F042457.

Zhu X, Stephens M (2017). “Bayesian large-scale multiple regression with summary statistics from genome-wide association studies.” The Annals of Applied Statistics, 11(3). doi:10.1214/17-aoas1046.

